# Combined EZH1/2 and DNMT inhibition overcomes transcriptional barriers imposed by drug-induced epigenetic plasticity in colorectal cancer cells

**DOI:** 10.1101/2025.09.16.676613

**Authors:** Alison A. Chomiak, Ashley K. Wiseman, Joel A. Hrit, Yanqing Liu, Stephanie Stransky, Ying Cui, Xiangqian Kong, Michael J. Topper, Stephen B. Baylin, Simone Sidoli, Rochelle L. Tiedemann, Scott B. Rothbart

## Abstract

Abnormal DNA methylation patterning is a defining epigenetic hallmark of human cancers and is therapeutically targetable with DNA methyltransferase inhibitors (DNMTi’s). However, epigenetic plasticity involving Polycomb Repressive Complex 2 (PRC2)-dependent H3K27 methylation can constrain transcriptional responses to DNMTi’s and may contribute to their limited activity in solid tumors. Here, we compare pharmacologic strategies targeting EZH1 and EZH2 in combination with DNMT inhibition in colorectal cancer (CRC) cells and define the molecular features that distinguish EZH2-selective from dual EZH1/2 inhibition. We find that dual EZH1/2 inhibition produces more robust transcriptional activation when combined with DNMT inhibition and more extensive depletion of H3K27 methylation states than EZH2-selective inhibition. Genetic and pharmacologic analyses implicate EZH1-dependent H3K27me1 as an adaptive chromatin barrier to PRC2 inhibitor-mediated chromatin remodeling. Dual EZH1/2 inhibition overcomes this barrier, increasing chromatin accessibility and H3K27 acetylation (H3K27ac) at previously silenced regulatory regions. EZH2 inhibition also induces a non-canonical bivalent chromatin state in which persistent DNA methylation maintains transcriptional repression despite acquisition of the activating H3K27ac mark. Combined DNMT inhibition resolves this additional drug-induced chromatin barrier, enhancing the activation of tumor-suppressive and stress response-associated transcriptional programs. Collectively, our studies reveal adaptive chromatin barriers that limit EZH inhibitor-mediated transcriptional activation and demonstrate how combined EZH1/2 and DNMT inhibition overcomes these consequences of drug-induced epigenetic plasticity in CRC cells. These findings provide a mechanistic framework for further evaluation of combined DNMT and EZH1/2 inhibition in solid tumors.

## Introduction

Abnormal DNA methylation patterning and dysregulated histone post-translational modification (PTM) signaling are defining epigenetic hallmarks of human cancer that drive tumor initiation and progression through their effects on transcriptional reprogramming (1,2). Unlike genetic mutations, epigenetic alterations can be reversed through targeting the enzymes that write, erase, and read these marks (3). The most clinically advanced class of drugs targeting epigenetic regulators are nucleoside analog DNA methyltransferase inhibitors (DNMTi’s) like azacitidine (vidaza; AZA) and 5-aza-2’-deoxycytidine (decitabine; DAC). These drugs induce DNA hypomethylation and restore the expression of epigenetically silenced tumor suppressor genes (TSG’s) and transposable elements (TE’s) that stimulate innate immune signaling through the induction of viral mimicry (4,5). Although DNMTi’s have achieved clinical success with U.S. Food and Drug Administration (FDA) approvals for several hematological malignancies (6–8), their efficacy in solid tumors has remained limited, in part because of pharmacologic constraints that include short half-lives in circulation and dose-limiting toxicities that restrict clinically achievable exposures (9,10).

In addition to these pharmacologic barriers, the molecular responses to DNMTi’s are constrained by epigenetic plasticity. Our group and others have shown that DNMTi-induced DNA hypomethylation promotes compensatory chromatin remodeling that counteracts transcriptional reactivation through the acquisition or redistribution of repressive histone PTMs (11–14). Polycomb Repressive Complex 2 (PRC2) is a prominent mediator of this response. PRC2 catalyzes mono-, di- and tri-methylation of lysine 27 on histone H3 (H3K27me1, H3K27me2, and H3K27me3), with H3K27me3 serving as a defining histone PTM signature of facultative heterochromatin. In our prior work, we showed that inhibition of PRC2-dependent H3K27me3 enhanced DNMTi-mediated transcriptional activation, viral mimicry, and anti-tumor responses (11,12), establishing PRC2-dependent chromatin regulation as a central mediator of DNMTi-induced epigenetic plasticity and a targetable pathway to limit adaptive resistance to DNMTi therapy.

The catalytic activity of PRC2 is mediated by the paralogous methyltransferases EZH1 and EZH2, which have overlapping but distinct biochemical and biological functions (15–18) EZH2 is generally considered the dominant PRC2 catalytic subunit in proliferating cells and has consequently received the greatest attention as a therapeutic target in cancer (19,20). These efforts have generated multiple small-molecule inhibitors that differ substantially in their relative selectivity for EZH2 versus EZH1 (21–24). However, whether EZH1 and EZH2 make distinct contributions to the adaptive chromatin response to DNMT inhibition remains poorly understood. Here, we address this question by comparing molecular effects of EZH2-selective and dual EZH1/2 inhibition, alone and in combination with DNMT inhibition, in colorectal cancer (CRC) cells. Our findings reveal that pharmacologic targeting of PRC2 catalytic activity generates distinct chromatin states that constrain transcriptional activation, including EZH1-dependent H3K27me1 retention and the acquisition of H3K27ac at persistently DNA-methylated regulatory regions. By comparing these chromatin states induced by EZH2-selective versus dual EZH1/2 inhibitors, we establish a mechanistic basis for the enhanced transcriptional activity of combined EZH1/2 and DNMT inhibition. Together, these studies highlight the importance of PRC2 inhibitor selectivity and adaptive chromatin remodeling in the rational design of epigenetic combination therapies.

## Results

### Dual EZH1/2 inhibition is more effective than EZH2-selective inhibition at enhancing DNMTi-induced transcription

Our prior studies showed that the EZH2-selective inhibitor tazemetostat (TAZ) enhances DNMTi-induced transcriptional activation in CRC cells (11). Building on this work, we sought to compare the efficacy of different pharmacologic strategies targeting PRC2 in enhancing the re-expression of DNA methylation-silenced genes. To facilitate this, we leveraged our previously described *SFRP1*-Nanoluciferase (NLuc) transcriptional reporter line engineered in the HCT116 colon cancer cell line (11). In this system, promoter CpG island hypermethylation silences an endogenous allele of the TSG *SFRP1*, and insertion of NLuc into exon 2 of this locus provides a quantitative readout of *SFRP1* re-expression from its native chromatin context. Notably, this reporter was engineered in a *DNMT1* hypomorphic background with 20% lower global DNA methylation compared to parental cells (25), sensitizing the system to detect additional chromatin-dependent transcriptional responses.

We screened a diverse panel of PRC2-directed compounds, including experimental and clinically applied SAM-competitive inhibitors of EZH1 and EZH2, protein-protein interaction (PPI) disruptors targeting EED, and PROTAC degraders of PRC2 subunits (**Figure 1A-B** and **S1A-D**). Among these compounds, three SAM-competitive inhibitors – tulmimetostat (CPI-0209; TUL) (26,27), valemetostat (DS3201b; VAL) (28–30), and mevrometostat (PF-1497; MEV) (31,32) – were most effective at inducing re-expression of *SFRP1* when combined with DAC at a concentration (30 nM) that has limited single agent efficacy at inducing expression of this reporter locus (11). Consistent with our previous findings (11), PRC2 inhibitors that most robustly induced transcription exhibited minimal toxicity as single agents (**Figure S1C-D**), suggesting that several of these compounds have off-target toxicities. Notably, VAL, TUL, and MEV (the three top-performing inhibitors in this screen) are also among the most clinically advanced PRC2 inhibitors (27,29–31,33), suggesting that they share a therapeutically relevant mode of action.

**Figure 1.**
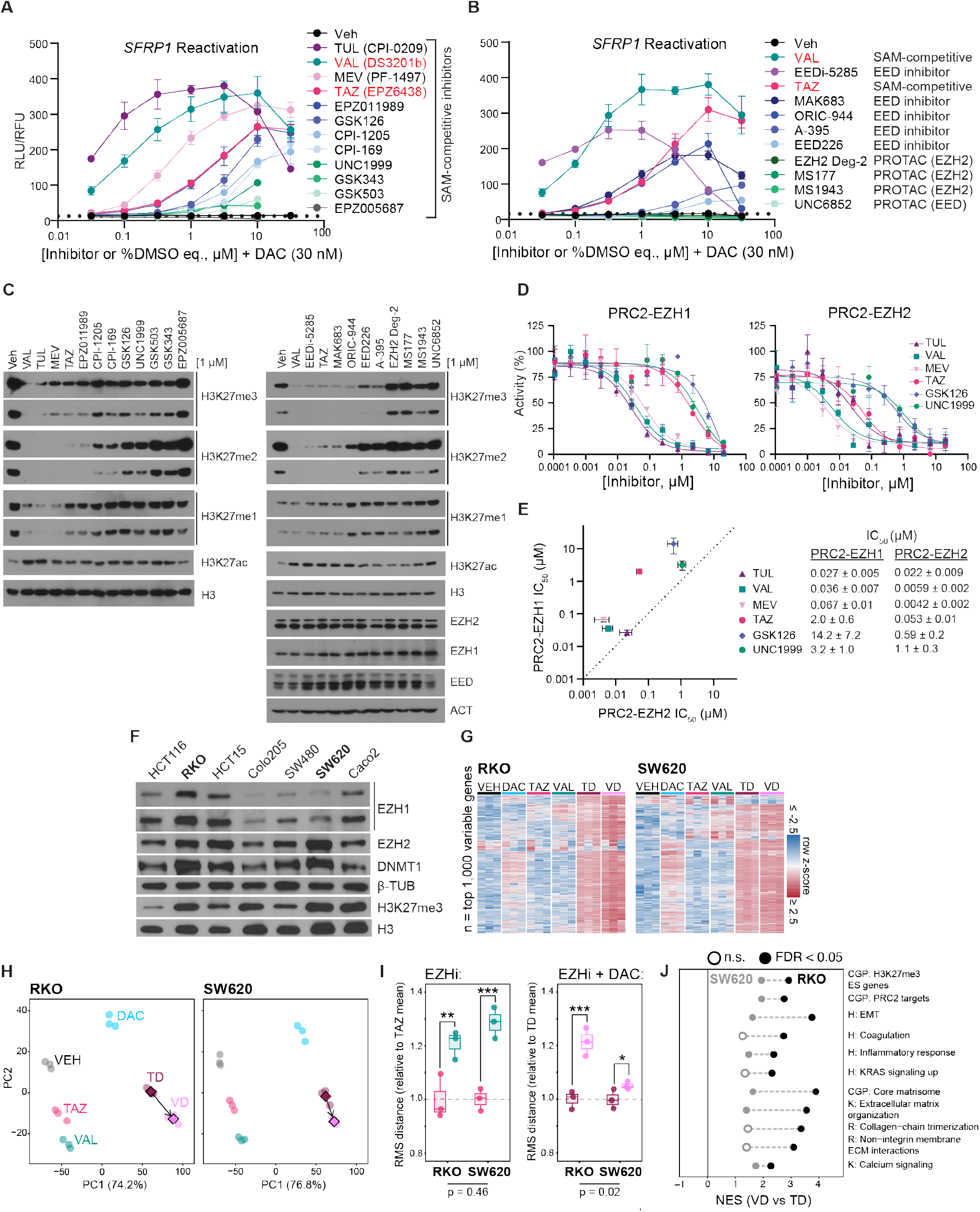
Dual EZH1/2 inhibition is more effective than EZH2-selective inhibition at enhancing DNMTi-induced transcription. **A-B)** NLuc reporter activity from the endogenous *SFRP1-*NLuc locus in HCT116 DNMT1 hypomorphic cells following 72-hour treatment with the indicated PRC2 antagonists titrated with a fixed concentration of DAC (30 nM). Red text indicates SAM-competitive inhibitors shared between experiments. Relative luminescence units (RLU) are normalized to relative fluorescence units (RFU) from multiplexed CellTiter-Fluor viability assays. Data are mean ± SD of technical triplicates and representative of n=3 biological replicates. Dotted black lines denote the maximal signal measured without the addition of DAC (see Figure S1A-B). TUL, tulmimetostat; VAL, valemetostat; MEV, mevrometostat; and TAZ, tazemetostat. **C)** Western blot analysis of the indicated proteins and PTMs from wild-type HCT116 cells following 72-hour treatment with vehicle (DMSO, % equivalent) or the indicated PRC2 inhibitors (1 µM) ordered approximately by ability to induce *SFRP1* expression in **A-B**. Short and long exposures are shown for H3K27 methylation states. **D)** *In vitro* methyltransferase activity assays using recombinant mononucleosome substrates and PRC2 core complexes (SUZ12, EED, and RbAp46/48) containing either EZH2 or EZH1 as the catalytic subunit. Selected SAM-competitive inhibitors from **A** were titrated as indicated. Normalized results are presented as percent PRC2 activity relative to no-inhibitor control, and data are the mean ± SD of technical triplicates. **E)** Comparison of IC_50_ values from **D**. Dotted line, x=y. **F)** EZH1, EZH2, DNMT1, H3K27me3, and corresponding loading controls were assessed across colorectal cancer cell lines by immunoblotting. RKO and SW620 were selected as models with relatively high and low EZH1 protein abundance, respectively. **G)** Unsupervised heatmaps of the top 1,000 most-variable protein-coding genes shared between RKO and SW620. Genes were ranked by the mean of their within-cell-line variances, thereby preventing either cell line from independently determining the feature space. Expression was row-standardized by z-score within each cell line and displayed using a common gene order. **H)** Principal-component analysis (PCA) using the same top 1,000 genes. Points represent biological replicates, diamonds denote TD and VD treatment centroids, and arrows indicate displacement from TD to VD. **I)** Transcriptome-wide treatment displacement was quantified as the root-mean-square (RMS) distance of each sample from the corresponding cell-line-specific VEH centroid. VAL increased transcriptional displacement relative to TAZ similarly in RKO and SW620 (cell-line × treatment interaction, (pval=0.458)). In contrast, VD produced a greater relative displacement from TD in RKO than in SW620 (interaction, pval=0.0195). Boxes show the interquartile range, center lines indicate medians, whiskers extend to 1.5 times the interquartile range, and points represent biological replicates. Interaction statistics were obtained from linear models containing cell line, treatment, and their interaction. **J)** Selected pathways enriched among genes differentially responsive to VD versus TD. Normalized enrichment scores from preranked gene-set enrichment analysis are shown for RKO and SW620. Filled points indicate FDR <0.05; open points indicate nonsignificant enrichment. **See also Figure S1.**

To define the mechanistic features distinguishing these compounds from less efficacious PRC2 inhibitors, we first analyzed global H3K27 methylation states by western blotting. The most effective compounds (TUL, VAL, and MEV) in our expression screen were also the most effective at reducing all three methylation states on H3K27 (**Figure 1C** and **S1E-F**). Notably, the strongest distinction was observed for H3K27me1, whose depletion correlated more closely with SFRP1 re-expression than loss of H3K27me2 or H3K27me3.

Prior *in vitro* biochemical studies, genetic knockout experiments, and differentiation studies in mouse embryonic stem cells have implicated EZH1 in maintaining H3K27me1, particularly when EZH2 is reduced or absent (15–17,34,35). Because both VAL and TUL have been reported to inhibit EZH1 in addition to EZH2 (26,28), we hypothesized that dual EZH1 and EZH2 inhibition contributes to their enhanced transcriptional activation potential compared to other inhibitors screened. Consistent with this hypothesis, TUL, VAL, and MEV were potent inhibitors of both EZH1- and EZH2-containing PRC2 complex methyltransferase activity towards recombinant nucleosome substrates compared to less efficacious compounds in our screen (**Figure 1D-E**).

To further investigate the potential contribution of EZH1 inhibition to this enhanced transcriptional response, we reasoned that the advantage of a dual EZH1/2 inhibitor over an EZH2-selective inhibitor would be evident in cells with elevated EZH1 levels. We therefore surveyed EZH1 protein expression across a panel of CRC cell lines and selected EZH1-high RKO and EZH1-low SW620 cells for comparative transcriptome analysis (**Figure 1F)**. Cells were treated with the EZH2-selective inhibitor TAZ or the dual EZH1/2 inhibitor VAL, alone or in combination with DAC, at concentrations that produced comparable reductions in H3K27me3 within each cell line (**Figure S1G**). Consistent with their relative protein abundance, EZH1 transcript levels were higher in RKO than in SW620 cells and were not substantially altered by treatment, whereas EZH2 transcript abundance was similar across the evaluated conditions (**Figure S1H**).

To compare global transcriptional responses, we performed unsupervised hierarchical clustering using the 1,000 protein-coding genes with the greatest mean within-cell-line variance across the two cell lines. Treatment-dependent transcriptional differences were more pronounced in RKO cells, particularly following combined VAL and DAC (VD) treatment, whereas transcriptional profiles were more similar across treatment groups in SW620 cells (**Figure 1G**). Principal-component analysis further revealed greater separation between TAZ plus DAC (TD) and VD treatment in RKO cells than in SW620 cells (**Figure 1H**).

We next quantified transcriptional displacement as the root-mean-square (RMS) distance of each treated sample from its respective cell-line-specific VEH centroid. VAL produced greater transcriptional displacement than TAZ in both cell lines, with no significant difference in the relative single-agent response between RKO and SW620 cells (**Figure 1I**). In contrast, the additional transcriptional displacement induced by VD relative to TD was significantly greater in EZH1-high RKO cells than in EZH1-low SW620 cells (**Figure 1I**). This relationship remained directionally consistent when the analysis was repeated using the 2,000, 5,000, or 10,000 most-variable genes (**Figure S1I**), indicating that the result was not dependent on the selected feature-space threshold.

To characterize the transcriptional programs preferentially activated by dual EZH1/2 inhibition, we performed gene set enrichment analysis comparing VD and TD treatments. In RKO cells, genes preferentially induced by VD relative to TD were enriched for PRC2- and H3K27me3-associated gene sets, as well as pathways related to epithelial-mesenchymal transition, extracellular matrix organization, inflammatory signaling, KRAS signaling, and calcium signaling (**Figure 1J**). These enrichments were generally weaker or absent in SW620 cells, consistent with the more pronounced transcriptional responses to VD in the EZH1-high background.

Collectively, these results identify dual EZH1/2 inhibition and depletion of all three H3K27 methylation states as distinguishing features of PRC2 inhibitors that most effectively enhance DNMTi-mediated transcriptional activation. The greater transcriptional response of dual EZH1/2 inhibition relative to EZH2-selective inhibition in EZH1-high CRC cells further supports a contribution of EZH1 inhibition to this enhanced activity. Together, these findings suggest that simultaneous targeting of EZH1 and EZH2 provides a more effective strategy for overcoming PRC2-mediated constraints on DNMTi-induced transcription than selective EZH2 inhibition.

### EZH1 maintains H3K27me1 following genetic or pharmacologic inhibition of EZH2

To better understand the role of EZH1 in regulating H3K27 methylation in CRC cells and its contribution to the differential effects of EZH2-selective versus dual EZH1/2 inhibition, we generated isogenic EZH1- and EZH2-knockout (KO) RKO cell lines using CRISPR/Cas9, alongside control-KO lines targeting a gene desert on chromosome 12. RKO cells were chosen due to their high endogenous expression levels of both EZH1 and EZH2 (**Figure 1F**). Successful gene disruption in clonally expanded cell populations was confirmed by western blot analyses and genomic sequencing (**Figure 2A** and **S2A**). Western blot analysis of all three H3K27 methylation states in these KO lines revealed that EZH1-KO had minimal effects on global H3K27 methylation levels, whereas EZH2-KO showed reduced global levels of H3K27me2 and H3K27me3, accompanied by a marked increase in H3K27me1 levels relative to control-KO lines (**Figure 2B**). These effects were observed both in the absence and presence of DAC. To validate these findings in an antibody-independent manner, and to more comprehensively assess broader histone PTM remodeling effects of these genetic deletions, we performed quantitative histone PTM mass spectrometry on these KO lines. Consistent with western blot analyses, EZH1-KO cells showed minimal changes in the distribution of H3K27 methylation states compared to control-KO cells (**Figure 2C**). In contrast, EZH2-KO cells showed pronounced reductions in H3K27me2 and H3K27me3 on both canonical H3.1 and variant H3.3, coupled with increased H3K27me1 on H3.1 and maintenance of H3K27me1 on H3.3 (**Figure 2C**). EZH2 loss was also associated with increased H3K36 methylation, consistent with the reported antagonistic relationship between H3K27 and H3K36 methylation (36,37), along with several additional minor PTM changes compared to both control-KO and EZH1-KO clones (**Figure S2B**).

**Figure 2.**
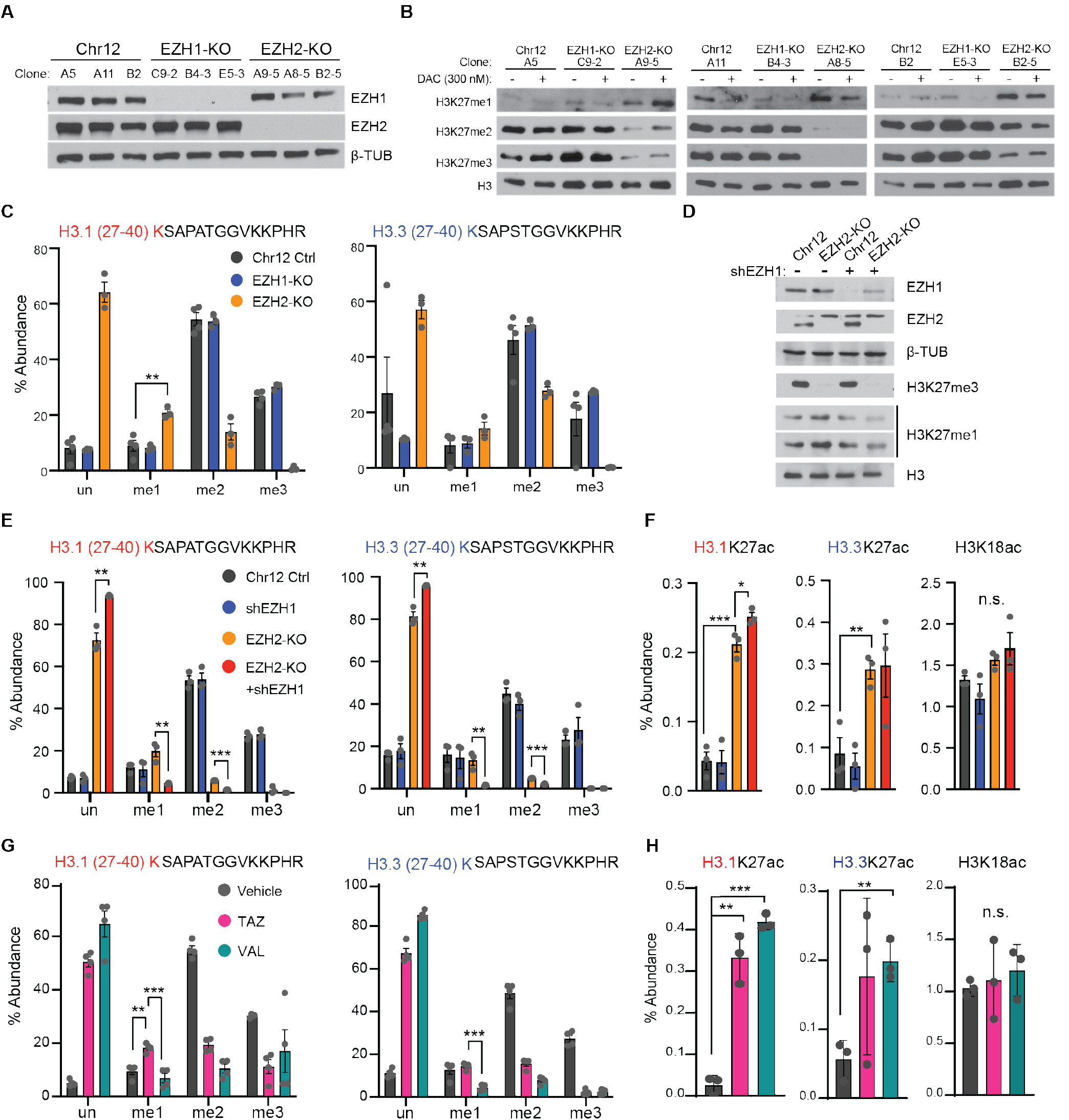
EZH1 maintains H3K27me1 following genetic or pharmacologic inhibition of EZH2. **A)** Western blot validation of EZH1 and EZH2 protein expression in RKO CRISPR/Cas9 KO clones targeting EZH1, EZH2, or a gene-desert control region on chromosome 12 (Chr12). Alphanumeric labels denote independently derived clonal populations. **B)** Western blot analysis of global H3K27 methylation states in RKO CRISPR KO clones treated with vehicle or DAC (300 nM) for 72 hours. **C)** Histone PTM mass spectrometry analysis of relative abundances for H3K27 methylation states on H3.1 and H3.3 in individual RKO CRISPR clones. Data are mean ± SD of three clones per genotype. **D)** Western blot analysis of the indicated proteins in RKO Chr12 control (clone B2) and EZH2-KO (clone A8-5) cells following shEZH1 induction with doxycycline for 72 hours. **E-F)** Histone PTM mass spectrometry analysis of relative abundances of H3K27 methylation states on H3.1 and H3.3 (**E**) or the indicated acetylated H3 peptides (**F**) in RKO Chr12 control and EZH2-KO cells following shEZH1 induction. Data are mean ± SD of n=3 biological replicates. **G-H)** Histone PTM mass spectrometry analysis of relative abundances of H3K27 methylation states on H3.1 and H3.3 (**G**) or the indicated acetylated H3 peptides (**H**) in RKO cells treated with the indicated EZH inhibitors (1 µM) for 72 hours. Data are mean ± SD of n=4 biological replicates. Red or blue “K” denotes modified lysine residues. Statistical significance was calculated using multiple unpaired t-tests. n.s., not significant; *p<0.05, **p<0.01, ***p<0.001. **See also Figure S2.**

To determine whether EZH1 contributes to the elevated H3K27me1 observed following EZH2 loss, we introduced a doxycycline-inducible EZH1-targeting shRNA into EZH2-KO cells. This approach enabled acute EZH1 depletion (KD) while circumventing the lethality associated with combined EZH1-and EZH2-KO in these cells, which is likely a consequence of targeting both catalytic and non-catalytic functions of EZH1/2 (15,38,39). EZH1 KD attenuated the EZH2-KO-associated increase in H3K27me1, as demonstrated by both western blotting and histone PTM mass spectrometry (**Figure 2D-E)**. Dual EZH1/2 depletion also produced additional gains in H3K36 methylation compared to EZH2-KO alone (**Figure S2C).** Notably, both EZH2-KO cells and dual EZH1/2-depleted cells showed increased H3K27 acetylation (H3K27ac), but not H3K18ac (**Figure 2F**). Given that both residues are substrates for p300/CBP histone acetyltransferase activity, these data suggest selective remodeling of H3K27ac following dual EZH1/2 depletion.

We next asked whether pharmacologic inhibition of EZH1 and EZH2 recapitulates these genetic phenotypes. We treated RKO cells with either the EZH2-selective inhibitor TAZ or the dual EZH1/2 inhibitor VAL at 1 µM, a concentration at which both compounds comparably inhibit EZH2 catalytic activity *in vitro*, and where VAL shows greater EZH1 inhibition (**Figure 1D-E**). Consistent with EZH2 genetic ablation, treatment of RKO cells with TAZ, alone or in combination with DAC, reduced H3K27me2 and H3K27me3 while increasing H3K27me1 (**Figure 2G** and **S2D-E**). In contrast, treatment with VAL, alone or in combination with DAC, resulted in loss of all three H3K27 methylation states, phenocopying results of combined EZH1-KD and EZH2-KO (**Figure 2G** and **S2D-E**). Similar to genetic perturbation, both TAZ and VAL increased H3K27ac levels without affecting H3K18ac or other histone acetylation sites (**Figure 2H** and **S2F**).

Collectively, these genetic and pharmacologic studies show that EZH1 contributes to the maintenance of H3K27me1 following EZH2 loss or selective inhibition. Importantly, dual EZH1/2 inhibition overcomes this residual H3K27me1 retention, providing a mechanistic distinction from EZH2-selective inhibition that may contribute to its greater efficacy in enhancing DNMTi-induced transcription.

### EZH1-dependent H3K27me1 redistribution following selective EZH2 inhibition is associated with restricted chromatin accessibility

Having established that EZH1 sustains H3K27me1 following selective EZH2 inhibition, we next mapped this response across the genome to identify the affected chromatin contexts and determine its relationship to chromatin accessibility. We first evaluated the specificity and performance of commercially available H3K27me1 antibodies (along with all other histone PTM antibodies used for chromatin immunoprecipitation (ChIP) in this study) using histone peptide microarrays and measurements of DNA mass enriched by ChIP (**Figure S3A-E**). We identified one H3K27me1 antibody with robust epitope specificity, consistent chromatin mass capture across antibody titrations, and treatment-dependent changes concordant with histone PTM mass spectrometry (**Figure S3A** and **S2E**). In contrast, other commercial H3K27me1 antibodies exhibited off-target recognition of mono-methylated epitopes and/or substantial lot-to-lot variability (**Figure S3B-C**).

Using this validated antibody, we performed sans spike-in quantitative chromatin immunoprecipitation sequencing (siQ-ChIP-seq) (40,41) to quantify genome-wide H3K27me1 distributions in vehicle (VEH)-, TAZ-, and VAL-treated RKO cells. Integration of these data with H3K27me3 ChIP-seq (12) and whole genome bisulfite sequencing (WGBS) data for DNA methylation (42) from the same cell line revealed that H3K27me1 was enriched across broad, highly DNA methylated domains that were depleted of H3K27me3 (**Figure 3A**). Chromatin Hidden Markov Model (ChromHMM) analysis showed that H3K27me1 was predominantly enriched across actively transcribed gene bodies and genic enhancers (active genic ChromHMM annotations) in RKO cells (**Figure 3B**), a pattern that is consistent with reported H3K27me1 localization in mouse embryonic and human hematopoietic stem cells (43,44). In contrast, H3K27me3 distributions were enriched across homogenously marked Polycomb-associated regions (ReprPCWk, ReprPC, TssBiv, EnhBiv), coinciding with reduced DNA methylation and H3K27me1 levels (**Figure 3A-B**).

**Figure 3.**
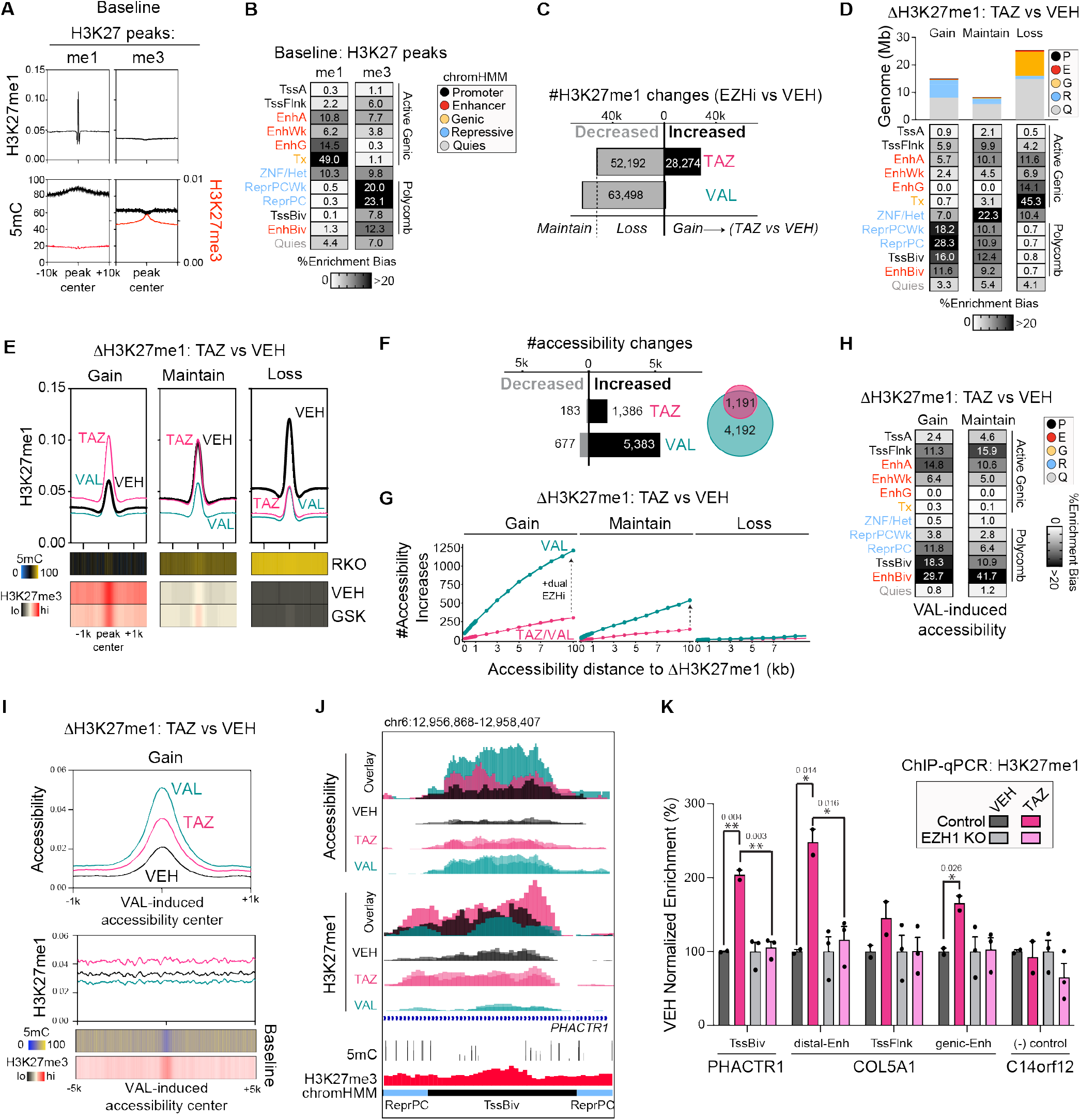
EZH1-dependent H3K27me1 redistribution following selective EZH2 inhibition is associated with restricted chromatin accessibility. **A)** Baseline relationships among H3K27me1, H3K27me3, and DNA methylation in RKO cells. Aggregate profiles are centered on baseline H3K27me1 or H3K27me3 peaks and display the corresponding histone PTM signal together with DNA methylation (5mC; black) and H3K27me3 (red) across the indicated windows. **B)** ChromHMM state enrichment of baseline H3K27me1 and H3K27me3 peaks. Values report enrichment bias relative to the genomic representation of each chromatin state, normalized by all states represented within the queried regions. States are grouped as active genic, Polycomb-associated, or quiescent and colored by their broader functional annotation. **C)** Numbers of genomic regions exhibiting decreased or increased H3K27me1 following selective EZH2 inhibition with Tazemetostat (TAZ) or dual EZH1/2 inhibition with Valemetostat (VAL), relative to vehicle (VEH). **D)** Genomic coverage and ChromHMM-state enrichment of three H3K27me1 response classes: TAZ-specific H3K27me1 gain (“Gain”), regions maintained following TAZ but lost following VAL (“Maintain”), and regions exhibiting H3K27me1 loss following both treatments (“Loss”). Stacked bars show total genomic coverage, and tables report enrichment bias across ChromHMM states. **E)** Aggregate H3K27me1 profiles for VEH-, TAZ-, and VAL-treated cells across the Gain, Maintain, and Loss classes. Heatmaps below show baseline DNA methylation and H3K27me3 signal across the same regions, centered on the corresponding H3K27me1 peaks. **F)** Numbers of regions exhibiting decreased or increased chromatin accessibility following TAZ or VAL treatment. The proportional Venn diagram summarizes the overlap between regions with increased accessibility following the two treatments. **G)** Cumulative numbers of TAZ- and VAL-induced accessibility increases occurring in direct overlap with, or within the indicated edge-to-edge distance of, Gain, Maintain, or Loss H3K27me1 regions. Dashed arrows highlight the additional accessibility response produced by dual EZH1/2 inhibition. **H)** ChromHMM-state enrichment of VAL-induced accessibility increases associated with Gain or Maintain H3K27me1 regions. Values report enrichment bias relative to the genomic representation of each state. **I)** Aggregate accessibility and H3K27me1 profiles centered on VAL-induced accessibility increases associated with TAZ-specific H3K27me1 gain. Heatmaps show baseline DNA methylation and H3K27me3 surrounding the same accessibility sites. **J)** Representative genome-browser view at the *PHACTR1* locus showing accessibility and H3K27me1 signal following VEH, TAZ, or VAL treatment, together with baseline DNA methylation, H3K27me3, and ChromHMM annotation. Overlay tracks display the combined treatment profiles. **K)** H3K27me1 ChIP–qPCR validation at the indicated ChromHMM annotated regions in control and EZH1-knockout RKO cells treated with VEH or TAZ. Points represent independent replicates, bars summarize normalized enrichment, and statistical comparisons and p-values are indicated above the corresponding brackets. Abbreviations: 5mC, 5-methylcytosine, DNA methylation; TssBiv, bivalent transcription start site; TssFlnk, transcription start-site flanking region; Enh, enhancer; ReprPC, Polycomb-repressed chromatin; Tx, transcribed gene bodies; ZNF/Het, Zinc Finger clusters/Heterochromatin; Quies, Quiescent chromatin. **See also Figures S3,4.**

Changes in H3K27me1 distribution were distinct following treatment with EZH2-selective (TAZ) versus dual EZH1/2 (VAL) inhibitors. Consistent with our prior analyses, VAL treatment globally reduced H3K27me1, whereas TAZ treatment induced both gains and losses (**Figure 3C**). We binned these treatment-dependent responses into three groups according to their H3K27me1 response to TAZ. “Gain” regions gained H3K27me1 with TAZ but showed no detectable change with VAL. “Maintain” regions retained H3K27me1 with TAZ but lost it with VAL. “Loss” regions lost H3K27me1 with both inhibitors (**Figure 3C**). Notably, all regions that lost H3K27me1 with TAZ also lost it with VAL, whereas additional VAL-dependent losses were captured in the “Maintain” group.

These three response groups differed in their baseline H3K27me1/me3 profiles, DNA methylation levels, and chromatin states. “Gain” regions had low baseline H3K27me1, high H3K27me3, and low DNA methylation levels and were enriched within repressive Polycomb (ReprPC, ReprPCWk), bivalent enhancer (EnhBiv), and bivalent promoter (TssBiv) chromatin states (**Figure 3D-E**). These data suggest that following EZH2 inhibition and the accompanying loss of H3K27me2/me3, residual EZH1 activity promotes H3K27me1 within these canonical PRC2 domains. Supporting this interpretation, TAZ-induced gains in H3K27me1 occurred at previously identified PRC2 target loci (**Figure S4A**). Gain regions were also enriched for Mef2, Hox, and Fox transcription factor binding motifs that have previously been associated with EZH1 activity in the contexts of cardiac regeneration and hematopoietic stem cell differentiation (**Figure S4B**) (17,18,45,46).

“Maintain” regions had intermediate baseline levels of H3K27me1, H3K27me3, and DNA methylation (**Figure 3D-E**), suggestive of heterogenous epigenetic patterning across the RKO cell population. These regions encompassed both PRC2-associated states and broader repressive ChromHMM annotations that included ZNF/Repeat and Heterochromatin regions (**Figure 3D-E**). Notably, these regions demonstrated an “epigenetic switch” between DNA methylation and H3K27me3 when DNMT1 was inhibited with the DNMT1 inhibitor GSK3484862 (**Figure 3E**), suggesting that they represent regulatory regions particularly sensitive to combined disruption of DNA methylation and PRC2 activity.

“Loss” regions, in contrast, were characterized by high baseline H3K27me1, high DNA methylation, and low H3K27me3 (**Figure 3E**). These regions mapped predominantly to actively transcribed gene bodies, with a smaller fraction mapping to enhancer elements, and overlapped extensively with H3K27me1-enriched domains in VEH-treated cells (**Figure 3B, D**). Comparable reductions in H3K27me1 following TAZ or VAL treatment (**Figure 3E**) suggest these regions of mono-methylation are regulated by EZH2 rather than EZH1.

To directly relate DNA methylation levels to EZH inhibitor-induced H3K27me1 gains and losses, we performed enzymatic methyl-sequencing (EM-seq) (47) on DNA fragments enriched by H3K27me1 ChIP (me1EM-seq). Consistent with conclusions made from integrative analysis of H3K27me1 ChIP-seq and WGBS datasets (**Figure 3A** and **3E**), me1EM-seq confirmed that H3K27me1-enriched regions were densely methylated and reflected DNA methylation patterns associated with the distinct classes of EZH inhibitor-induced H3K27me1 responses described above (**Figure S4C**). Importantly, DNA methylation levels were unchanged by EZH inhibitor monotherapy (**Figure S4D**), and DNA methylation patterns remained highly concordant between TAZ and VAL treatments across regions of H3K27me1 redistribution (**Figure S4E**). In contrast, DNMTi treatment reduced DNA methylation across all regions of H3K27me1 redistribution (**Figure S4F**). Collectively, these data demonstrate that H3K27me1, like DNA methylation, marks both repressive and active regions of the genome and undergoes context-dependent remodeling in response to EZH2-selective versus dual EZH1/2 inhibition.

We next performed ATAC-seq to determine whether these EZH inhibitor-induced H3K27me1 dynamics were associated with changes in chromatin accessibility. Notably, VAL treatment increased chromatin accessibility to a greater extent than TAZ (**Figure 3F**), and more than 30% of regions gaining accessibility with VAL mapped within 10kb of an H3K27me1 Gain or Maintain region (**Figure 3G**). In contrast, H3K27me1 Loss regions were not associated with increased chromatin accessibility following either treatment (**Figure 3G**). VAL-induced chromatin accessibility gains associated with H3K27me1 redistribution preferentially localized to bivalent promoters and enhancers (TssBiv, EnhBiv) embedded within Polycomb repressive domains (**Figure 3H-I**). These findings associate the persistence or acquisition of H3K27me1 following selective EZH2 inhibition to a chromatin accessibility barrier that is not observed with dual EZH1/2 inhibition.

Finally, to determine whether this adaptive H3K27me1 response required EZH1, we selected several regulatory regions where VAL-induced chromatin accessibility gains were adjacent to regions of H3K27me1 redistribution (**Figure 3J, S4G**). We then measured H3K27me1 by ChIP-qPCR in control-and EZH1-KO CRISPR clones (**Figure 2A, 3K**). Consistent with ChIP-seq results, TAZ increased H3K27me1 at each tested region in control cells, whereas these gains were absent in EZH1-KO cells (**Figure 3K**). These data validate that the TAZ-induced H3K27me1 response at these regulatory elements requires EZH1.

Collectively, these findings reveal context-dependent and non-redundant contributions of EZH1 and EZH2 to H3K27me1 patterning. EZH2 contributes to H3K27me1 within active, DNA methylated genomic regions, whereas EZH1 supports H3K27me1 deposition or maintenance within Polycomb-regulated chromatin following EZH2 inhibition. The association of this EZH1-dependent H3K27me1 response within regulatory elements that become accessible upon dual EZH1/2 inhibition further supports a model in which adaptive H3K27me1 remodeling contributes to the maintenance of a repressive chromatin state following selective EZH2 inhibition.

### EZH2 inhibition induces a non-canonical H3K27ac/DNA methylation bivalent chromatin state that limits transcriptional activation

Although TAZ and VAL monotherapies produced modest transcriptional responses, their combination with DAC markedly enhanced transcriptional activation (**Figure 1G**). We therefore sought to determine how DNMT inhibition alters the chromatin response to EZH inhibition to promote cooperative gene activation. ATAC-seq analysis revealed that DAC alone had marginal effects on chromatin accessibility at a concentration that enhanced transcription when combined with EZH inhibition, whereas its combination with either TAZ (TD) or VAL (VD) substantially increased chromatin accessibility (**Figure 4A**). To distinguish the genomic responses to these combinations, we used overlap analysis to classify regions gaining chromatin accessibility according to their response to monotherapy and combination treatment (**Figure S5A**). Regions selectively opened by VD or exhibiting greater than 20% increased accessibility relative to TD were classified as “VD-selective,” whereas regions showing comparable accessibility gains with both combinations were classified as “Comparable” (**Figure 4B, S5B**). Nearly 50% of VD-selective chromatin accessibility gains occurred within 10kb of regions that gained or maintained H3K27me1 following selective EZH2 inhibition (**Figure 4C**), linking the adaptive H3K27me1 barrier described above to regulatory regions preferentially opened by combined EZH1/2 and DNMT inhibition. Notably, VD-selective accessibility gains were enriched at bivalent promoters (TssBiv), whereas accessibility gains shared by TD and VD were more broadly distributed across bivalent enhancers (**Figure 4D**).

**Figure 4.**
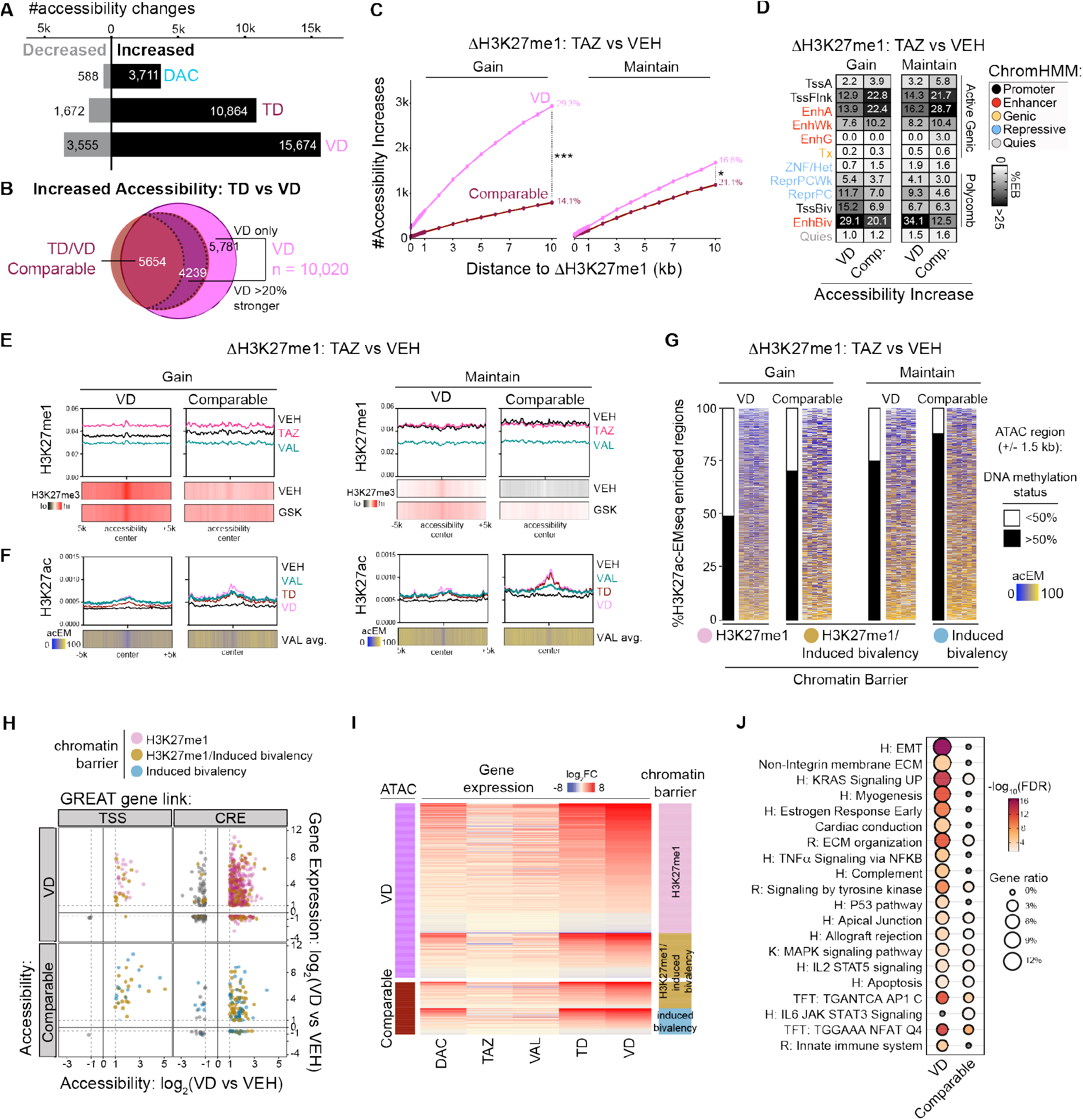
EZH2 inhibition induces a non-canonical H3K27ac/DNA methylation bivalent chromatin state that limits transcriptional activation. **A)** Numbers of regions exhibiting decreased or increased chromatin accessibility following decitabine (DAC), tazemetostat plus decitabine (TD), or valemetostat plus decitabine (VD), relative to VEH. **B)** Proportional overlap of TD- and VD-induced accessibility increases. Shared regions were classified as “Comparable” when the TD and VD responses were similar or as “VD >=20% stronger” when accessibility increased by more than 20% in VD relative to TD. Regions detected only following VD were classified as “VD only.” VD accessibility comprises the VD-only and VD >= 20% stronger classes. **C)** Cumulative numbers of VD and Comparable accessibility increases occurring in direct overlap with, or within the indicated distance of, TAZ-specific H3K27me1 Gain or TAZ-maintained H3K27me1 regions. Percentages indicate the fraction of each accessibility class occurring within 10 kb. The indicated comparison was evaluated using a two-sided Fisher’s exact test; *** p< e-100, *p<e-10. **D)** ChromHMM state enrichment of VD and Comparable accessibility increases associated with H3K27me1 Gain or Maintain regions. Values report enrichment bias relative to the genomic representation of each chromatin state, normalized by all states represented within the queried regions. **E)** Aggregate H3K27me1 profiles following VEH, TAZ, or VAL treatment across VD and Comparable accessibility regions associated with H3K27me1 Gain or Maintain. Heatmaps below show H3K27me3 signal following VEH or GSK treatment across the same regions. **F)** Aggregate H3K27ac profiles following VEH, VAL, TD, or VD treatment across the same accessibility and H3K27me1 response classes. Heatmaps show the mean methylation of H3K27ac-associated DNA measured by VAL acEM-seq. **G)** Position-resolved VAL acEM-seq methylation across the indicated ATAC regions. Each row represents an individual accessibility region centered within a +/- 1.5 kb window and ranked from lowest to highest mean DNA methylation. Only regions with acEM-seq signal in at least six of twelve 250-bp bins were retained (500 bp bins presented for clarity). Adjacent binary bars classify sufficiently covered H3K27ac-associated fragments as methylated when their mean VAL acEM-seq methylation was >= 50% or below threshold when methylation was <50%. **H)** Relationship between VD-induced accessibility and gene expression for genes linked to accessibility regions by GREAT. Genes are separated by promoter-proximal transcription start-site (TSS) or distal candidate regulatory element (CRE) assignment and by VD or Comparable accessibility. Points are colored according to the associated chromatin-barrier class; gray points represent genes without an assigned barrier class. Dashed lines indicate the predefined accessibility and expression fold-change thresholds. **I)** RNA-expression changes across DAC, TAZ, VAL, TD, and VD treatments for GREAT-linked genes associated with VD or Comparable accessibility. Genes are ordered within each accessibility class and annotated by the associated chromatin-barrier state. **J)** Gene-set enrichment among genes linked to VD or Comparable accessibility. Dot size represents the gene ratio, and color indicates -log_10_ false-discovery rate (FDR). H, Hallmark; R, Reactome; K, KEGG; TFT, transcription-factor target gene set. **See also Figure S5.**

To further define the chromatin contexts associated with these differential accessibility responses, we integrated EZH inhibitor-induced H3K27me1 dynamics with baseline and DNMT1i-induced H3K27me3 distributions (**Figure 4E**). VD-selective accessibility gains were preferentially associated with regions that gained or maintained H3K27me1 following TAZ treatment (**Figure 4E**), consistent with residual EZH1 activity constraining chromatin opening following selective EZH2 inhibition. Notably, these regions separated into two distinct epigenetic contexts. Regions with the strongest TAZ-induced H3K27me1 gains had high baseline H3K27me3 that remained largely unchanged following DNMT1 inhibition, whereas regions that maintained H3K27me1 following TAZ exhibited increased H3K27me3 following DNMT1 inhibition (**Figure 4E**). These findings suggest that dual EZH1/2 inhibition more effectively overcomes both pre-existing Polycomb-mediated repression and compensatory H3K27 methylation associated with DNMT inhibition.

Having linked EZH1-dependent H3K27me1 retention to restricted chromatin accessibility, we next asked whether depletion of H3K27 methylation permits acquisition of activating histone modifications at these regulatory regions. Our global histone PTM mass spectrometry analysis showed increased H3K27ac following EZH inhibition (**Figure 2H**). Consistent with this observation, VAL monotherapy increased H3K27ac across regions preferentially opened by VD (**Figure 4F**). However, despite acquiring H3K27ac, these regions exhibited limited chromatin opening with VAL alone, and VAL monotherapy produced only modest transcriptional activation (**Figure 3F, 1A,G**). Notably, the addition of DAC further increased H3K27ac relative to VAL monotherapy, with VD producing greater H3K27ac enrichment than TD (**Figure 4F**). These findings suggested that H3K27ac acquisition alone was insufficient to promote robust chromatin opening and raised the possibility that persistent DNA methylation constitutes an additional repressive barrier at regulatory regions that acquire H3K27ac following EZH inhibition.

To test this hypothesis, we performed EM-seq on H3K27ac-enriched chromatin (acEM-seq), enabling direct measurement of DNA methylation within H3K27ac-associated DNA fragments (**Figure S5C**). Strikingly, H3K27ac acquired following VAL treatment frequently co-occurred with methylated DNA (**Figure 5G**), revealing a drug-induced chromatin state characterized by the co-existence of an activating histone PTM (H3K27ac) and a repressive DNA modification (DNA methylation). We refer to this as a non-canonical drug-induced bivalent chromatin state. Importantly, the persistence of DNA methylation within H3K27ac-marked chromatin provides a potential explanation for the limited transcriptional response to EZH inhibitor monotherapy despite acquisition of an activating histone PTM. Together, these findings identify two distinct adaptive chromatin features that may constrain transcriptional activation in EZH1-dependent H3K27me1 retention following selective EZH2 inhibition and H3K27ac/DNA methylation-associated non-canonical bivalency following EZH inhibition.

**Figure 5.**
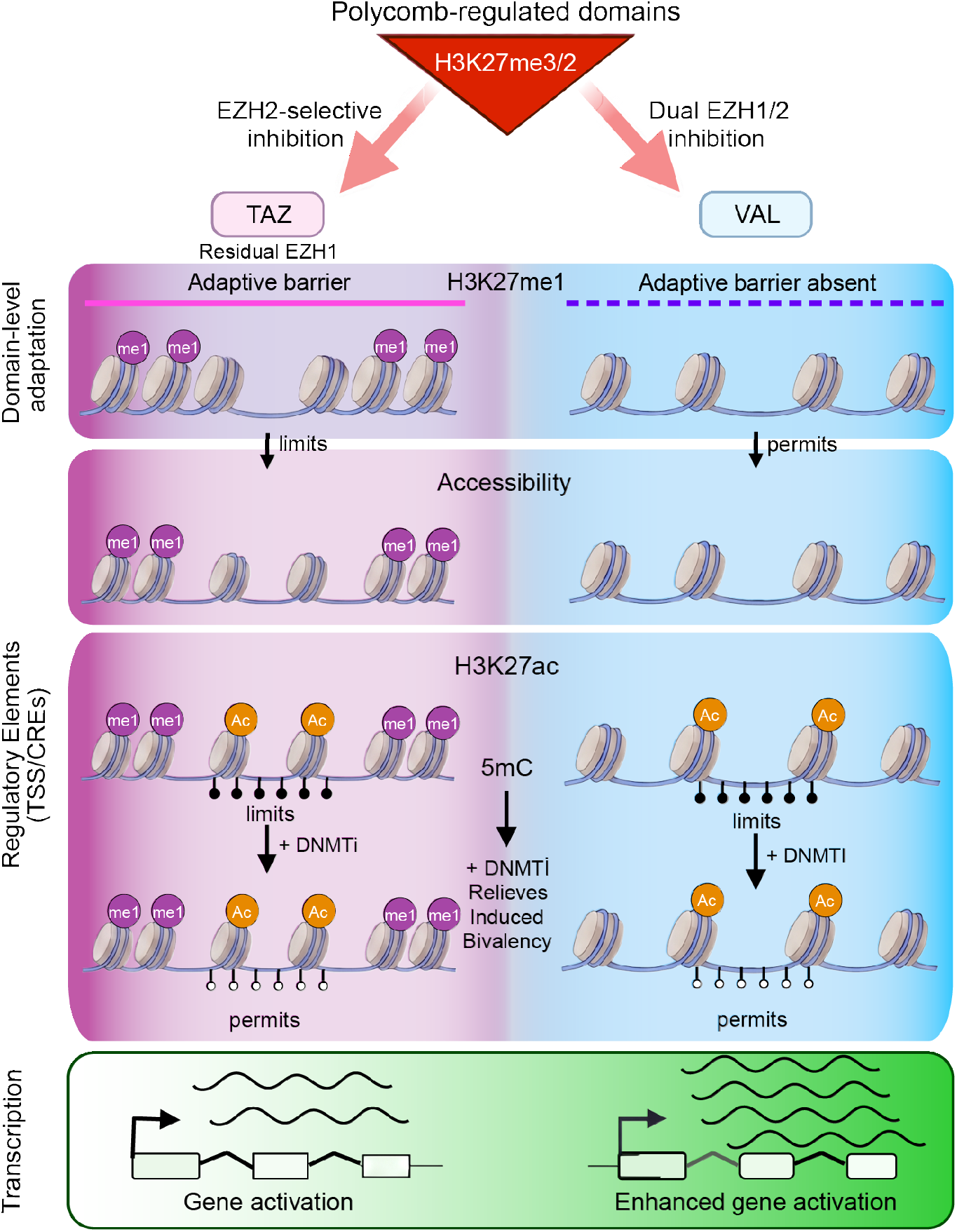
Summary of key results. Combination EZH and DNMT inhibition overcomes drug-induced chromatin barriers to transcriptional activation in colorectal cancer cells. EZH2-selective inhibition (left), in combination with DNMT inhibition, resolves a therapy-induced bivalent H3K27ac-5mC chromatin state, permitting transcriptional activation. However, residual EZH1 activity maintains an adaptive H3K27me1 barrier that limits transcriptional responses. Dual EZH1/2 inhibition (right), in combination with DNMT inhibition, overcomes both the therapy-induced bivalent chromatin state and the EZH1-dependent H3K27me1 barrier, enabling enhanced chromatin accessibility and transcriptional activation relative to EZH2-selective inhibition.

Finally, we sought to determine the relationship between these drug-induced chromatin features and transcriptional activation. We integrated chromatin accessibility and gene expression data using the Genomic Regions Enrichment of Annotations Tool (GREAT) (48) to associate “VD-selective” and “Comparable” chromatin accessibility gains at cis-regulatory elements (CREs) and transcription start sites (TSSs) with expression of their putative target genes (**Figure 4H**). This analysis revealed distinct relationships between chromatin remodeling and transcriptional responses. VD-selective accessibility gains associated with adaptive H3K27me1 retention following TAZ treatment were strongly linked to genes preferentially activated by VD, whereas accessibility gains shared by TD and VD were predominantly associated with regions exhibiting EZH inhibitor-induced non-canonical bivalency (**Figure 4H-I**). These findings support a model in which EZH1-dependent H3K27me1 retention preferentially constrains transcriptional activation following selective EZH2 inhibition, while persistent DNA methylation at H3K27ac-marked regulatory elements represents an additional barrier shared by both EZH inhibitor strategies. Importantly, VD treatment enhanced the activation of transcriptional programs commonly suppressed in cancer, linking the ability to overcome these chromatin constraints with the re-expression of potentially tumor-suppressive pathways (**Figure 4J**).

Collectively, these findings support a model in which EZH inhibition induces a partially permissive chromatin state characterized by H3K27ac acquisition in the context of persistent DNA methylation, priming previously silenced regulatory regions for transcriptional activation (**Figure 5**). Subsequent DNMT inhibition relieves this constraint, promoting chromatin accessibility and transcriptional activation at associated genes. However, selective EZH2 inhibition leaves an additional barrier in the form of EZH1-dependent H3K27me1 retention or redistribution, particularly within Polycomb-regulated promoters and enhancers. By more effectively depleting all three H3K27 methylation states, dual EZH1/2 inhibition overcomes this adaptive response and permits more extensive chromatin remodeling when combined with DNMT inhibition (**Figure 5**). Together, these findings establish a mechanistic framework in which combined EZH1/2 and DNMT inhibition overcomes distinct, drug-induced chromatin constraints to enhance transcriptional activation of cancer-relevant gene programs.

## Discussion

We previously showed that EZH2 inhibition enhances the transcriptional activating effects of DNMTi in CRC cells by limiting compensatory PRC2-dependent chromatin repression (11). Here, we extend these findings by showing that dual EZH1/2 inhibition is more effective than EZH2-selective inhibition at enhancing DNMTi-induced transcription. Through integrative pharmacologic, genetic, and epigenomic analyses, we identify EZH1-dependent H3K27me1 retention or redistribution and the acquisition of H3K27ac at persistently DNA methylated genomic regions as adaptive chromatin responses to EZH2 inhibition. Our findings support a model in which these drug-induced chromatin states represent distinct barriers to transcriptional activation that can be overcome through combined inhibition of DNMTs and both catalytic subunits of PRC2. More broadly, these results highlight how drug-induced epigenetic plasticity can influence therapeutic responses and provide a mechanistic rationale for selecting epigenetic inhibitor combinations that target compensatory chromatin regulatory pathways.

A key finding of this study is that PRC2 inhibitors differ substantially in their ability to enhance DNMTi-mediated transcriptional activation despite sharing the ability to suppress H3K27me3. Among the compounds screened, the most effective inhibitors were SAM-competitive molecules that potently inhibited both EZH1- and EZH2-containing PRC2 complexes and promoted depletion of all three H3K27 methylation states. Notably, these compounds include several of the most clinically advanced PRC2 inhibitors, suggesting that dual EZH1/2 inhibition may be a key pharmacologic contributor to their therapeutic success. Although differences in biochemical potency, cellular target engagement, and other compound-specific properties may also influence their efficacy, our findings identify the ability to inhibit H3K27me1 as a distinguishing feature of dual EZH1/2 inhibition. These observations suggest that suppression of H3K27me3 alone may be insufficient to predict transcriptional consequences of PRC2 inhibition, particularly in the context of combination treatment with DNMTi.

Our genetic and pharmacologic studies identify EZH1 as an important regulator of H3K27me1 following disruption of EZH2 catalytic activity. Consistent with the established dominance of EZH2 in proliferating cells (49), EZH1 depletion alone had minimal effects on global H3K27 methylation states in RKO cells. In contrast, genetic deletion or selective pharmacologic inhibition of EZH2 substantially reduced H3K27me2 and H3K27me3 while maintaining or increasing H3K27me1. Importantly, genetic depletion of EZH1 in an EZH2-deficient background attenuated this H3K27me1 response, phenocopying the effects of dual EZH1/2 inhibition. These findings are consistent with previous studies in mouse embryonic stem cells implicating EZH1 in H3K27me1 maintenance when EZH2 activity is reduced or absent (16,17,34,35) and extend this relationship to cancer cells undergoing pharmacologic PRC2 inhibition. Because EZH1 and EZH2 also possess functions independent of their catalytic activities (15,38,39), the concordance between conclusions drawn from genetic and pharmacologic perturbations is particularly informative. Together, these results support a catalytic contribution of EZH1 to H3K27me1 maintenance following EZH2 inhibition.

Genome-wide profiling further revealed that EHZ1- and EZH2-dependent H3K27me1 regulation is strongly influenced by chromatin context. At baseline, H3K27me1 was broadly enriched within actively transcribed, DNA methylated gene bodies where both EZH2-selective and dual EZH1/2 inhibitors reduced its abundance. In contrast, selective EZH2 inhibition promoted H3K27me1 accumulation within distinct Polycomb-annotated regions marked by H3K27me3 at baseline, including bivalent promoters and enhancers. These regions were preferentially associated with chromatin accessibility gains following dual EZH1/2 inhibition, suggesting that residual EZH1 activity contributes to the maintenance of a restrictive chromatin environment when EZH2 activity is selectively inhibited. Notably, these findings distinguish the established association of H3K27me1 with transcriptionally active chromatin from its adaptive redistribution into repressive Polycomb-regulated domains following EZH2 inhibition (43). Although the precise contribution of H3K27me1 to chromatin repression remains to be determined, our findings support a model in which EZH1-dependent H3K27me1 remodeling is associated with an adaptive chromatin barrier that limits the response to EZH2-selective inhibition.

A second key conceptual advance from this work is the identification of a non-canonical, drug-induced bivalent chromatin state characterized by the coexistence of H3K27ac and DNA methylation. Although EZH2 inhibition promoted H3K27ac accumulation at previously repressed regulatory regions, these changes were accompanied by limited chromatin accessibility gains and transcriptional activation. Using acEM-seq, we showed that H3K27ac acquired following EZH2 inhibition frequently remained associated with methylated DNA, and that this persistent DNA methylated state maintains a repressed chromatin state despite the presence of this transcription-associated histone PTM. Importantly, this chromatin configuration is mechanistically distinct from canonical H3K4me3/H3K27me3 bivalency and illustrates how epigenetic modifications generally associated with opposing transcriptional states can function to resist epigenetic plasticity induced by pharmacologic perturbation of chromatin modifiers.

These findings, together with our prior work (11), provide a mechanistic framework for understanding the enhanced transcriptional responses to combined DNMT and EZH1/2 inhibition. DNMT inhibitor-induced DNA hypomethylation promotes compensatory H3K27me3 to maintain a repressive chromatin environment at select genomic regions, whereas EZH2 inhibition induces an EZH1- and H3K27me1-associated chromatin accessibility barrier at a subset of Polycomb-repressed loci and promotes H3K27ac accumulation across genomic regions that maintain their repressed state by DNA methylation. Combined DNMT and dual EZH1/2 inhibition targets these multi-layered mechanisms of compensatory repression, producing greater chromatin accessibility and transcriptional activation than either monotherapy or combined DNMT and EZH2-selective inhibition. As pathways of transcriptional activation associated with combined DNMT and EZH1/2 inhibition included gene sets involved in inflammatory signaling, cellular stress responses, and other tumor suppressive processes, our findings suggest that the efficacy of epigenetic combination therapies depends not only on reversing pre-existing repressive chromatin states but also on preventing or overcoming drug-induced adaptive chromatin responses.

Our mechanistic studies have potential translational implications for the application of epigenetic therapy combinations. The enhanced transcriptional response to dual EZH1/2 inhibition in EZH1-high RKO cells compared to EZH1-low SW620 cells suggests that EZH1 abundance may influence the relative benefit of dual versus EZH2-selective inhibitors. However, additional studies will be required to determine whether EZH1 expression independently predicts treatment response across broader genetic and tumor contexts. Moreover, the greater molecular efficacy of dual EZH1/2 inhibitors in our experimental systems does not necessarily establish their therapeutic superiority, which will also depend on compound-specific pharmacology, tolerability, and effects on non-malignant cells. Nevertheless, our findings identify PRC2 inhibitor selectivity and adaptive H3K27 modifications as important considerations for optimizing DNMTi-based combination strategies. More broadly, this work establishes a framework for understanding how the coordinated targeting of DNA and PRC2-associated methylation can overcome drug-induced chromatin constraints and enhance the reactivation of epigenetically silenced transcriptional program in CRC cells.

### Limitations of the Study

Several limitations of this study warrant consideration. First, although our studies support a contribution of EZH1 to an H3K27me1-associated chromatin barrier and the establishment of a non-canonical bivalent chromatin state following EZH2 inhibition, these conclusions are based on correlations made from integrative and orthogonal measurements of chromatin modifications, chromatin accessibility, and transcription following genetic and pharmacologic perturbations. Second, the precise role of H3K27me1 in restricting chromatin accessibility and transcriptional activation, and the mechanism by which DNA methylation dominates over H3K27ac in regions of drug-induced non-canonical bivalency remains unresolved. Third, our studies were conducted in colorectal cancer cell lines, and genomic analyses were conducted in bulk RKO cell populations. Therefore, it remains to be determined the extent to which the identified chromatin responses vary across individual cells and broader tumor contexts. Furthermore, although the differential responses to EZH1-high and EZH1-low cell lines suggest that EZH1 abundance may influence sensitivity to dual EZH1/2 inhibitors, its predictive value remains to be established across genetically diverse models. Finally, these experiments were conducted in two-dimensional culture systems and therefore do not address how these epigenetic combinations influence tumor growth, therapeutic response, or interactions with the tumor microenvironment. Future studies using organoid, patient-derived, and *in vivo* models will be required to establish the broader biological and translational relevance of these findings.

## Resource Availability

### Lead Contact

Requests for further information and resources should be directed to and will be fulfilled by the lead contact, Dr. Scott Rothbart.

### Materials Availability

Plasmids and cell lines generated by the study are available upon request from the lead contact.

### Data and Code Availability

All sequencing and EPIC array data are available on the Gene Expression Omnibus (GEO) under the following accession numbers and will available upon publication: ChIP-seq (GSE320400), EM-seq (GSE320401), EPIC-array (GSE320533), RNA-seq (GSE320534), and ATAC-seq (under curation – GEO accession code will be made available prior to publication). Final code used for bioinformatic analysis is deposited on our GitHub page (https://github.com/rleetied/EZH1_EZH2_study) and archived on Zenodo (10.5281/zenodo.24086061).

## Acknowledgements

We thank members of the Rothbart laboratory and the Epigenetic Therapies SPORE for helpful discussions. We also thank Amy Johnson, Ibukunoluwa Sodiya, Hyoungjoo Lee, Colt Capan, and Molly Soper-Hopper for sample preparation, data acquisition, and analysis of proteomics data. Proteomics experiments were conducted in the Van Andel Institute (VAI) Mass Spectrometry Core (RRID:SCR_024903) with financial support from the VAI Metabolism and Nutrition (MeNu) Program (RRID:SCR_ 027494). We acknowledge support from the VAI Genomics Core (RRID:SCR_022913) and the VAI Bioinformatics and Biostatistics Core (RRID:SCR_024762). This work was supported by grants from the National Institutes of Health to S.B.R., M.J.T., and S.B.B. (P50CA254897), A.A.C. (F32CA225043), Y.L. (F99CA305570), R.L.T. (R50CA315355), and a grant from the American Cancer Society to S.B.R. (RSG-21-031-01-DMC).

## Author Contributions

Conceptualization: A.A.C, A.K.W., R.L.T., S.B.R.; Data Curation: A.A.C, A.K.W., J.A.H, Y.L., S. Stransky, R.L.T.; Formal Analysis: A.A.C, A.K.W., J.A.H, R.L.T.; Funding Acquisition: A.A.C, Y.L., M.J.T., R.L.T., S.B.B., S.B.R.; Investigation: A.A.C., A.K.W., J.A.H., Y.L., S. Stransky., R.L.T.; Methodology: A.A.C., A.K.W., J.A.H, S. Stransky., R.L.T.; Project Administration: S.B.R.; Resources: A.A.C., A.K.W., Y.C., X.K., R.L.T.; Software: R.L.T.; Supervision: R.L.T., S. Sidoli., S.B.R.; Validation: A.A.C., A.K.W., J.A.H., R.L.T.; Visualization: A.A.C., A.K.W., J.A.H., R.L.T.; Writing – original draft: A.A.C., A.K.W., R.L.T., S.B.R.; Writing – review & editing: all authors.

## Declaration of generative AI and AI-assisted technologies in the manuscript preparation process

During the preparation of this work, the authors used OpenAI Codex to assist with computational code development and troubleshooting, data visualization, and documentation of analytical workflows. All analyses, visualizations, and workflows were reviewed, verified, and edited by the authors as needed. The authors take full responsibility for the content of the published article.

## STAR Methods

### Cell culture

Caco2 (ATCC, HTB-37), Colo205 (ATCC, CCL-222), HCT15 (ATCC, CCL-225), HCT116 (ATCC, CCL-247), RKO (ATCC, CRL-2577), SW480 (ATCC, CCL-228), SW620 (ATCC, CCL-227) colon cancer cell lines and HEK293T (ATCC, CRL-3216) were purchased from American Type Culture Collection (ATCC) and cultured according to ATCC recommendations. These cells lines were incubated according to ATCC recommendations at 5% CO_2_ and 37°C in EMEM (ATCC, 20-2003), RPMI-1640 (ATCC, 30-2001), McCoy’s (Gibco, 16600-082), Leibovitz’s L-15 (ATCC, 30-2008), or DMEM (Gibco, 11995-065) culture media. RKO were maintained in RPMI 1640 (Gibco, 11875-093) culture media supplemented with L-glutamine (Gibco, 25030-081), 0.1 mM non-essential amino acids (Gibco, 11140-050), and 1 mM Sodium Pyruvate (Gibco, 11360070). All media were supplemented with 10% Fetal Bovine Serum (FBS, Millipore Sigma F0926) and 1% penicillin/streptomycin (Life Technologies, 15140-122). All cell lines were routinely tested for mycoplasma contamination using the Venor GeM Mycoplasma Detection Kit (Sigma, MP0025-1KT) according to the manufacturer’s instruction.

### CRISPR/Cas9-mediated knockout cell line generation

Cloning was performed as previously reported (50). Briefly, pSpCas9(BB)-2A-Puro (PX459) V2.0 (Addgene, plasmid #62988) was digested with restriction enzyme BbsI-HF (New England Biolabs, R3539L) in rCUTSMART buffer (New England Biolabs, B6004S) and purified using Monarch Spin DNA Gel Extraction and Monarch Spin PCR & DNA Cleanup Kits (New England Biolabs, T1120S and T1130S). Oligo pairs for genes of interest (see below) were annealed and ligated into the digested vector using T4 ligase (ThermoFisher, EL0014) and transformed into XL-10 Gold ultracompetent bacterial cells (Agilent, 200315). Miniprepped plasmids (Qiagen, 27104) were confirmed by whole plasmid with Oxford Nanopore sequencing (Plasmidsaurus).

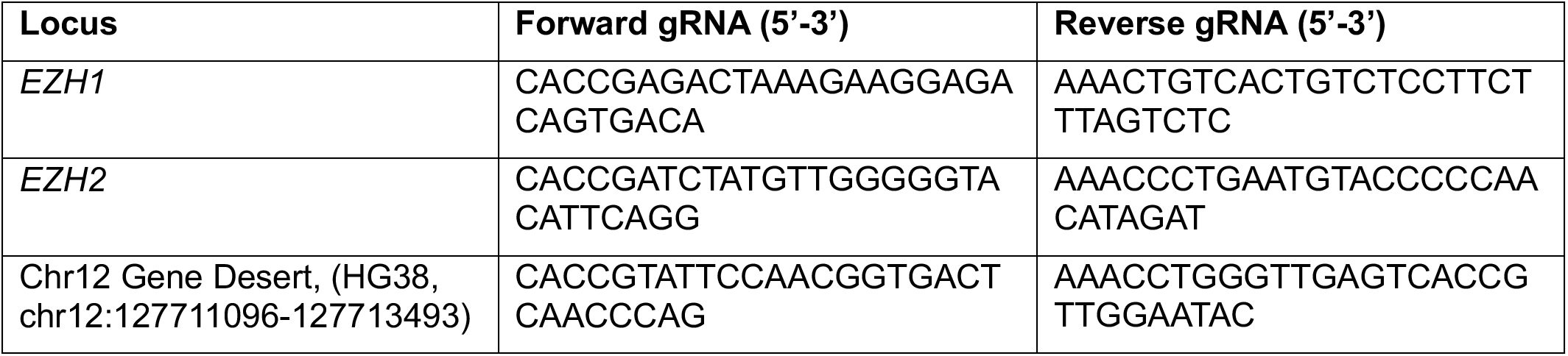

Plasmids were transfected into target cell lines at a 1 µg:2 µL ratio with XtremeGene HP (Sigma, 6366244001) and selected with 2 µg/mL puromycin (Sigma, P8833) for 48 hrs. Cells were split into single cell clonal populations through limiting dilution and individual knockout clones were confirmed by western blotting for the protein of interest and Sanger sequencing (Genewiz) across the CRISPR guide target region. To analyze mutations or indels on each allele for an individual clone, we first amplified a ∼400bp region around the CRISPR guide cut site (EZH2 For 5’-ACCATGCACAATATTTAGTTGGCT, EZH2 Rev 5’-TGATAGCACTCTCCAAGCTGC-3’, EZH1 For 5’-GTTTTGTTAAGCAGCTCTCATTGT-3’, EZH1 Rev 5’- CAGTGATCCTGGGTTTACCAGAG-3’), ligated the amplicon into a pGEM-T vector (Promega, A1360), and transformed the plasmid into JM109 cells (Promega, L2001) according to the manufacturer’s guidelines. Plasmid DNA was amplified directly from bacterial colonies in 96-well plates through rolling PCR amplification using TempliPhi according to the manufacturer’s protocol (GE Healthcare, 25-6400-10) and sent for Sanger sequencing (Genewiz).

### Inducible shRNA cell line generation

Inducible shRNA cell lines were generated as previously described (51,52). Briefly, the TET-pLKO-puro plasmid (Addgene, 21915) was digested with AgeI and EcoRI, and the fragment was gel purified. shRNA oligos targeting EZH1 were selected from the Broad Institute RNAi consortium shRNA library list. Oligos were ordered, annealed, and then ligated into the digested plasmid. Plasmid sequence was confirmed by Oxford Nanopore sequencing (Plasmidsaurus).

The plasmid for doxycycline-inducible shEZH1 knockdown was then transfected into HEK293T cells with pPAX2 (Addgene, 12260) and pMD2.G (Addgene, 12259) packaging plasmids to make lentivirus. RKO clones with CRISPR mutations for EZH2 (clone A8-5) or the gene desert Chr12 control (clone B2) were infected with virus media mixed with 1 mg/mL polybrene (Millipore, TR-1003-G) followed by selection with 2 µg/mL puromycin. To confirm knockdown, cells were treated with 20 ng/mL doxycycline (Cayman Chemical, 14422) for 72 hrs, collected, western blotted as below, and probed with an EZH1 antibody (Cell Signaling, 42088).

### Small molecule preparation and use

Small molecules were dissolved in 100% dimethyl sulfoxide (DMSO) at 5 or 10 µM and stored at -20°C. Cells were treated with each drug or vehicle (DMSO, % equivalent) for 72-hr treatment cycles with no media change. Drugs were obtained from the following sources: 5-aza-2’-deoxycytidine (DAC or decitabine; Sigma, A3656), A485 (Cayman Chemicals, 24119), A-395 (Cayman Chemicals, 20257), CPI-169 (Selleck Chemicals, S7616), CPI-1205 (lirametostat; Selleck Chemicals, S8353), EED226 (MedChemExpress, HY-101117), EEDi-5285 (MedChemExpress, HY-136977), EPZ0011989 (Selleck Chemicals, S7805), EPZ005687 (Cayman Chemical, 13966), EZH2 Degrader-2 (MedChemExpress, HY-157164), GSK126 (Selleck Chemicals, S7061), GSK343 (Selleck Chemicals, S7164), GSK503 (Cayman Chemical, 18531), MAK683 (MedChemExpress, S8983), mevrometostat (MEV or PF-1497; Chemietek, CT-PF0682), MS177 (Selleck Chemicals, E1163), MS1943 (MedChemExpress, HY-133129), ORIC-944 (Selleck Chemicals, E1967), tazemetostat (TAZ or EPZ6438; Selleck Chemicals, S7128), tulmimetostat (TUL or CPI-0209; Selleck Chemicals, E1497), UNC1999 (Caymen Chemical, 14621), valemetostat (VAL or DS-3210b; Chemietek, CT-DS3201), and UNC6852 (MedChemExpress, HY-130708).

### Cell viability and longitudinal proliferation assays

To measure cell viability, cells were plated in 96-well plates at a density of 3000-4000 cells/well and treated the following day. The CellTiter-Fluor kit (Promega, G6080) was used according to the manufacturer’s protocol, and fluorescence signal was measured using a BiotTek SynergyNeo2 Microplate Reader. Background signal from media-only wells was subtracted to obtain the final reported relative fluorescence units (RFU).

To measure RKO and SW620 cell longitudinal proliferation over time, cells were plated at a density of ∼1000-2000 cells/well (96-well plate) or ∼5000 cells/well (24-well plate) and allowed to adhere overnight. The following day, cells were treated with drugs in fresh media. Treated RKO cells were immediately placed in a Sartorius Incucyte S3 for imaging. For SW620, cells were replated and retreated a second time and directly placed in the Incucyte for imaging after an initial 72-hr treatment cycle.

To measure longitudinal proliferation of HCT116 following pretreatment, 100,000 cells/well were plated in a 6-well plate and allowed to adhere overnight. The following day, cells were treated with drugs in fresh media. After 72 hrs, cells were replated and retreated again (second treatment cycle). At six days (two 72-hr treatment cycles), 40,000 cells/well were plated into a 6-well plate and re-treated with drugs for a third and final time before being directly placed in the Incucyte for imaging.

For each well and treatment, the Incucyte captured 4 (96-well plate), 9-12 (24-well plate), or 12 (6-well plate) brightfield images per well at each timepoint. “Day 0” on each confluency graph represents the timepoint cells were placed into the Incucyte for image capture regardless of any preceding pretreatment cycles. The average confluency (%) per well and timepoint was calculated using the Sartorius Incucyte Live Cell Imaging and Analysis software v2020C (SW620 and RKO experiments) or v2024B (all HCT116 experiments and RKO experiments with the addition of A485).

### *SFRP1*-NanoLuciferase cell reporter assay

Engineering of the *SFRP1*-Nanoluciferase cell reporter cell line was previously reported (11). Briefly, a NanoLuciferase (NLuc) cassette was inserted into exon 2 of an endogenous *SFRP1* allele in DNMT1-hypomorph HCT116 colon cancer cells with reduced DNA methylation (25). To measure NLuc activity in *SFRP1*-NLuc reporter cells, the CellTiter-Fluor viability assay was duplexed with the Nano-Glo Luciferase Assay System (Promega, N1110). First, cells were plated and treated in 96-well plates as above for cell viability assays. To duplex the assays, CellTiter-Fluor was used at a 5x concentration preceding the Nano-Glo assay. NanoGlo reagent was applied according to the manufacturer’s protocol, and a BioTek SynergyNeo2 Microplate Reader was used to obtain relative luminescence units (RLU). After background subtraction from media-only wells, RLU were normalized to CellTiter-Fluor RFU to account for cell viability.

### siRNA transfection

ON-TARGETplus siRNA SMARTpools (5 µl of 20 µM stock) targeting EZH1 (Dharmacon, L-004217-00-0010) and non-targeting control (NTC) siRNA pools (Dharmacon, D-001810-10-05) or Lipofectamine RNAiMax (5 µL; ThermoFisher, 13778075) were each separately diluted into 100 µL Optimem (Life Technologies, 31985-062). After a 5 min separate incubation, diluted siRNA and RNAiMax were mixed together 1:1 and incubated for 20 min. McCoy’s Media supplemented with 10% FBS but no antibiotics was refreshed on attached cells that had been plated the previous day at 100,000/well in a 6-well plate. The siRNA/Lipid mixture was added dropwise to the fresh media for a final siRNA concentration of 50 nM. Cells were collected 72 hrs after transfection.

### Western blotting

Cells were lysed on ice in cold CSK lysis buffer as previously reported (11). 2-5 µg of total protein (to probe for histones) or 10-15 µg of total protein (for other protein targets) were size-separated by SDS-PAGE and transferred to PVDF membrane. Membranes were blocked for one hour at room temp (PBS, 0.1% Tween-20, and 5% BSA), washed in PBST (PBS supplemented with 0.1% Tween-20), and incubated in blocking buffer overnight at 4°C with the following antibodies: β-Actin (1:1,000; Cell Signaling Technology, 4970), β-Tubulin (1:50,000; Protein Tech, 66240), DNMT1 (1:1,000; Abcam, ab134148), EED (1:1,000; Cell Signaling Technology, 85322), EZH1 (1:1,000; Cell Signaling Technology, 42088), EZH2 (1:1,000; Cell Signaling Technology, 5246), H3 (1:50,000; EpiCypher, 13-0001), H3K27ac (1:1,000; Active Motif, 39133), H3K27me1 (1:1,000; Active Motif, 61015), H3K27me2 (1:2,000; Cell Signaling Technology, 9728), and H3K27me3 (1:2,000; Cell Signaling Technology, 9733). Histone antibodies were chosen based on specificity profiling with histone peptide microarrays as previously described (53). The following day, membranes were washed in PBST and incubated in HRP-conjugated secondary antibody (1:10,000; Sigma-Aldrich, GENA934) for 1 hour at room temp, washed again in PBST, incubated in Enhanced Chemiluminescence substrate (Cytiva, 29018903), and imaged with x-ray film.

### Recombinant nucleosome preparation

Recombinant mononucleosomes were reconstituted as previously described (54). Briefly, *Xenopus laevis* histones H2A, H2B, H3, and H4 were purchased from The Histone Source (Colorado State University). Lyophilized histones were resuspended in unfolding buffer (20 mM Tris-HCl pH 7.5, 7 M guanidine-HCl, and 10 mM DTT). Octamers were assembled by mixing histones H2A/H2B/H3/H4 and dialyzing into refolding buffer (10 mM Tris-HCl pH 7.5, 2 M NaCl, 1 mM EDTA, 5 mM BME) at 4°C, then purified by size exclusion chromatography using a Superdex 200 column (Cytiva). 175 bp DNA templates were generated by PCR amplification of the 147×601 Widom nucleosome positioning sequence and purified by ion exchange chromatography using a RESOURCE Q column (Cytiva). Octamers and DNA templates were mixed and assembled into nucleosomes via an 18-hour dialysis from RB high (10 mM Tris-HCl pH 7.5, 2 M KCl, 1 mM EDTA, and 1 mM DTT) to RB low (10 mM Tris-HCl pH 7.5, 250 mM KCl, 1 mM EDTA, and 1 mM DTT) at 4°C.

### *In vitro* PRC2 activity assays

Recombinant PRC2 complexes comprising SUZ12, EED, RbAp46, and RbAp48 with EZH1 or EZH2 were purchased (Active Motif, 31500 and 31387, respectively). PRC2 activity assays containing 20 nM PRC2 complex, 750 nM recombinant nucleosome substrate from above, 20 µM SAM, and inhibitor or vehicle (DMSO) were assembled in reaction buffer (20 mM Tris-HCl pH 8, 50 mM NaCl, 1 mM EDTA, 3 mM MgCl_2_, and 0.1 mg/mL BSA) in a final volume of 10 µL. Reactions were incubated at room temp overnight, then quenched by adding TFA to 0.5% v/v. PRC2 activity was measured using the MTase-Glo Methyltransferase Assay (Promega, V7601) according to the manufacturer’s instructions. IC_50_ values were generated by non-linear regression analysis using GraphPad Prism (RRID:SCR_002798).

### Sample preparation for histone PTM mass spectrometry

Histones were isolated by acid extraction as previously described (55). Briefly, cell pellets were lysed in hypotonic buffer (10 mM Tris–HCl pH 8.0, 1 mM KCl, 1.5 mM MgCl_2_, 1 mM DTT, 1 mM PMSF, 5 mM sodium butyrate, EDTA-free Roche mini protease inhibitor tablet (Sigma, 11836170001), and PhosSTOP Phosphatase Inhibitor tablet (Sigma, 4906837001) with rotation at 4°C for 30 min. Nuclei were then pelleted by centrifugation at 10,000 g for 10 min at 4°C, resuspended in cold 0.2 M H_2_SO_4_, rotated at 4°C for 2-4 hrs, and centrifuged at 16,000 g for 10 min at 4°C. Histones in the soluble fraction were precipitated by dropwise addition of cold trichloroacetic acid (33% v/v) followed by incubation with rotation at 4°C for at least 1 hr. Precipitated histones were pelleted by centrifugation at 16,000 g for 10 min at 4°C, washed with cold acetone in 0.1% HCl, pelleted again at 3,400 g for 5 min at 4°C, and washed again with cold acetone. A glass Pasteur pipette was used to gently wash precipitates that collected on the sides of tubes during spin steps. Samples were air-dried for 20 min on the benchtop before proceeding to chemical derivatization and digestion steps.

Acid extracted histones were propionylated by chemical derivatization as previously described (12,56) or with the following procedural modifications. Purified histones were resuspended in 100 mM ammonium bicarbonate (Sigma, 09830) with 20% acetonitrile (Fisher Chemical, LC/MS grade) at 25 µg histone/µL, propionylated with 5 µL ≥99% propionic anhydride (Sigma, 240311) immediately followed by the addition of 10 µL 20-22% ammonium hydroxide (Fisher, A470), pH was adjusted to ∼8.0 with additional propionic anhydride or ammonium hydroxide, and incubated for 10 min at 25°C. Propionylation was repeated once more before overnight tryptic digestion at a ratio of 50:1 histone:trypsin (Promega, V5073). After overnight digestion, samples were propionylated two more times as before and dried in a vacuum concentrator (Genevac, SpeedVac). Digests were cleaned using an Ultra-Micro C18 Spin Column (Harvard Apparatus, 74-7206), dried, and resuspended at a concentration of 1 µg/µL in 0.1% trifluoroacetic acid (TFA; Sigma, T6508) prior to mass spectrometry analysis.

### Data-independent acquisition (DIA) LC-MS/MS proteomics for histones and their PTMs

DIA analyses were performed on Orbitrap Exploris 480 coupled to Vanquish Neo system (Thermo Fisher Scientific). 1 μg of digested propionylated peptides were separated on a nano capillary liquid chromatography (LC) column (20 cm × 75 μm I.D., 365 μm O.D., 1.7 μm C18, CoAnn Technologies #HEB07502001718IWF) at a flow rate of 300 nL/min. Mobile phase A consisted of 0.1% formic acid (FA) in LC/MS-grade H_2_O (Fisher Scientific, LS118- 500) and mobile phase B consisted of 80% LC/MS-grade acetonitrile (Fisher Scientific, LS122500) and 0.1% FA in LC/MS-grade H_2_O. The LC gradient was: 3% B to 30% B over 51 min, 80% B over 5 min, 98% B over 2 min, and a hold at 98% B for 3 min, with a total gradient time of 60 min. Column temperature was kept constant at 45 °C using a customized column heater (Phoenix S&T). Full MS spectra were collected at 120,000 resolution (full width half-maximum; FWHM), and MS2 spectra at 30,000 resolution (FMWH). Both the standard automatic gain control (AGC) target and the automatic maximum injection time were selected. A precursor range of 300-1000 m/z was set for MS2 scans, and an isolation window of 30 m/z was chosen with a 1 m/z overlap for each scan cycle. 32% HCD collision energy was used for MS2 fragmentation. DIA data was processed in EpiProfile (version 2.1) (57) using the manufacturer’s default parameters.

### Sample Preparation for untargeted global proteomics

Samples were processed using the EasyPep Maxi MS Sample Prep Kit (Thermo Fisher Scientific, A45734) according to the manufacturer’s protocol. Briefly, up to 2X10^7^ cells were resuspended in 50 µL of lysis buffer and proteins were quantified using the Pierce BCA Protein Assay Kit (Thermo Fisher Scientific, 23227) following the manufacturer’s protocol. Plates were read at an absorbance of 562 nm using a Synergy LX Multi-Mode Reader, and Gen5 software was used for data analysis (BioTek/Agilent). Polynomial regression was used in the Gen5 software to calculate protein concentrations to a protein standard curve after an average blank absorbance subtraction. 25 µg of protein per sample was then reduced and alkylated at 95°C for 10 min, and samples were digested overnight with Trypsin/Lys-C at 30°C at a ratio of 10:1 (protein:enzyme (w/w)). Resulting peptides were cleaned with EasyPep Maxi kit peptide clean up columns and dried down in a vacuum concentrator. Samples were resuspended in 12.5 µL 0.1% FA (Fisher Scientific, LS118-1) and diluted 1:1 with 6 µL of 0.1% TFA (Fisher Scientific, LS119-500) in autosampler vials.

### Data-independent acquisition (DIA) LC-MS/MS for untargeted global proteomics

DIA analyses were performed on an Orbitrap Eclipse coupled to a Vanquish Neo system (Thermo Fisher Scientific) with a FAIMS Pro source (Thermo Fisher Scientific) located between the nanoESI source and the mass spectrometer.

Digested peptides (2 μg) were separated on a nano capillary column (20 cm × 75 μm I.D., 365 μm O.D., 1.7 μm C18, CoAnn Technologies #HEB07502001718IWF) at a flow rate of 300 nL/min. Mobile phase A consisted of 0.1% FA in LC/MS grade H_2_O (Fisher Scientific, LS118- 500) and mobile phase B consisted of 80% LC/MS grade acetonitrile (Fisher Scientific, LS122500) and 0.1% FA in in LC/MS grade H_2_O. The LC gradient was: 1% B to 24% B over 110 min, 85% B over 5 min, and 98% B over 5 min, with a total gradient time of 120 min.

The column temperature was kept constant at 50 °C using a customized column heater (Phoenix S&T). For FAIMS, the selected compensation voltage (CV) was applied (−40V, -55V, and -70V) throughout the LC-MS/MS runs. Full MS spectra were collected at 120,000 resolution (full width half-maximum; FWHM), and MS2 spectra at 30,000 resolution (FMWH). Both the standard automatic gain control (AGC) target and the automatic maximum injection time were selected. A precursor range of 380-985 m/z was set for MS2 scans, and an isolation window of 50 m/z was chosen with a 1 m/z overlap for each scan cycle. 32% HCD collision energy was used for MS2 fragmentation.

DIA data were processed in Spectronaut (version 18, Biognosys) using direct DIA. Data were searched against the *Homo sapiens* proteome. The manufacturer’s default parameters were used. Briefly, trypsin/P was set as the digestion enzyme, and two missed cleavages were allowed. Cysteine carbamidomethylation was set as a fixed modification, and methionine oxidation and protein N-terminus acetylation as variable modifications. Identification was performed using a 1% q-value cutoff on precursor and protein levels. Both peptide precursors and protein false discovery rate (FDR) were controlled at 1%. Ion chromatograms of fragment ions were used for quantification. For each targeted ion, the area under the curve between the XIC peak boundaries was calculated.

### Chromatin preparation

Cells were fixed in 1% formaldehyde in Dulbecco’s phosphate-buffered saline (DPBS) without calcium or magnesium (Gibco, 14190-144) 10 min at 25°C with shaking, quenched with 125 mM glycine for 5 min at room temp, scraped into cold DPBS, washed 2X with cold DPBS, flash frozen in liquid nitrogen, and stored at -80°C until use. Thawed pellets were lysed in 1 mL cell lysis buffer (5 mM PIPES pH 8.0, 85 mM KCl, 1% NP40, 100 mM PMSF, Roche EDTA-free protease inhibitor cocktail) for 20 min with rotation at 4°C and cleared by centrifugation at 2000 x g for 5 min at 4°C. Pelleted nuclei were washed in cold Micrococcal nuclease (Mnase) digestion buffer (50 mM Tris-HCl pH 8.0, 1 mM CaCl_2_, 300 mM sucrose, 100 mM PMSF, Roche EDTA-free protease inhibitor cocktail), pelleted by centrifuge, resuspended in 1 mL 37°C Mnase buffer with 50U Mnase (Worthington, LS004798), and incubated at 37°C shaking vigorously for 12 min. Digestion was stopped by addition of 10 mM EDTA and nuclei were transferred to a 1 mL milliTUBE (Covaris). Chromatin was sheared to a range of 300-600 base-pair fragments using a Covaris E220 evolution Focused ultrasonicator with the following parameters: Peak power (140.0), Duty Factor (5.0), Cycles/Burst (200), Duration (120 seconds), Temperature (4°C). Samples were centrifuged at 13,000 rpm at 4°C for 10 min and supernatants collected and stored at – 80°C.

To quantify chromatin, 20 µL of the above-digested chromatin was diluted in 80 µL elution buffer (50 mM NaHCO_3_, 1% SDS), 0.3 M NaCl, and 200 µg/mL proteinase K. Samples were incubated at 67°C overnight to reverse crosslinks and remove protein, followed by incubation with RNase A/T (Invitrogen EN0551) at 37°C for 30 min to remove RNA. DNA was purified using the Qiagen PCR purification kit (Qiagen, 28104) and quantified by Qubit dsDNA High Sensitivity kit (Invitrogen, Q32854). Correct DNA fragment size was confirmed by 2% agarose gel electrophoresis.

### Antibody validation for chromatin immunoprecipitation

Antibodies were analyzed for binding specificity using histone peptide arrays as previously described (53,58). Briefly, streptavidin-coated glass microscope slides that displayed a printed library of biotinylated singly- and combinatorially-modified (primarily methylation, acetylation, and phosphorylation) histone peptides were blocked with in hybridization buffer (1X PBS pH 7.6, 0.1% Tween, 5% BSA) followed by primary antibody hybridization. Following washing and hybridization of an Alexa Fluor 647 conjugated secondary antibody (1:5000; Life technologies, A21245), slides were scanned with an Innoscan 1100AL microarray scanner (Innopsys). Binding profiles were then analyzed using ArrayNinja (59) (**Figure S4**).

Chromatin enrichment analysis with antibody titrations was performed as previously described (41) with a fixed amounts of chromatin prepared as above (equivalent to 5 µg quantified DNA) and incubated with increasing amounts of antibody. Chromatin was immunoprecipitated with antibodies as below and total DNA mass quantified (**Figure S4**).

### Chromatin Immunoprecipitation (ChIP)

For immunoprecipitation, prepared chromatin (equivalent to 5 µg quantified DNA) was incubated with either 2.5 µL (2.5 µg) of H3K27ac antibody (ActiveMotif, 39133, lot 31521015) or 5 µL of H3K27me1 antibody (ActiveMotif, 61015, lot 03822017) overnight at 4°C with constant rotation. 5% input of normalized chromatin was removed and set aside. The next day, 30 µL of Dynabeads Protein G magnetic beads (ThermoFisher, 10004D) was added and samples were incubated for 3 hours at 4°C with rotation. Bead-immuno-chromatin complexes were then washed 3X for 5 min with rotation at room temperature with low salt WB1 (20 mM Tris-HCl pH 8.0, 150 mM NaCl, 2 mM EDTA, 1% Triton X-100, 0.1% SDS), 3X with high salt WB2 (20 mM Tris-HCl pH 8.0, 500 mM NaCl, 2 mM EDTA, 1% Triton X-100, 0.1% SDS), 1X with WB3 (20 mM Tris-HCl pH 8.0, 250 mM LiCl, 1 mM EDTA, 1% NP-40, 1% Na-deoxycholate) and 1X with TE (10 mM Tris-HCl pH 8.0, 1 mM EDTA pH 8.0). Beads were incubated in 100 µl of elution buffer shaking at 65°C for 30 min to elute immuno-chromatin complexes. Reverse crosslinking, protein digestion, RNA digestion, and DNA clean up were performed as above for both the IPs and the input samples to both quantify and isolate DNA for sequencing.

### ChIP-qPCR

ChIP-ed DNA was analyzed for enrichment by quantitative PCR (qPCR) using the KAPA SYBR FAST qPCR 2X master mix (KAPA biosystems KK4602). 5% of each total immunoprecipitated mass was put into each qPCR reaction and amplification was performed with BioRad CFX Real-time PCR detection system with an initial denaturation at 95 °C for 3 min, followed by 40 cycles of denaturation at 95 °C for 3 s and annealing/extension at 60 °C with the primers indicated in the table below for 20 s. Fluorescence was acquired at the end of each annealing/extension step. Each IP was normalized to its input, then each genomic region was normalized to Vehicle.

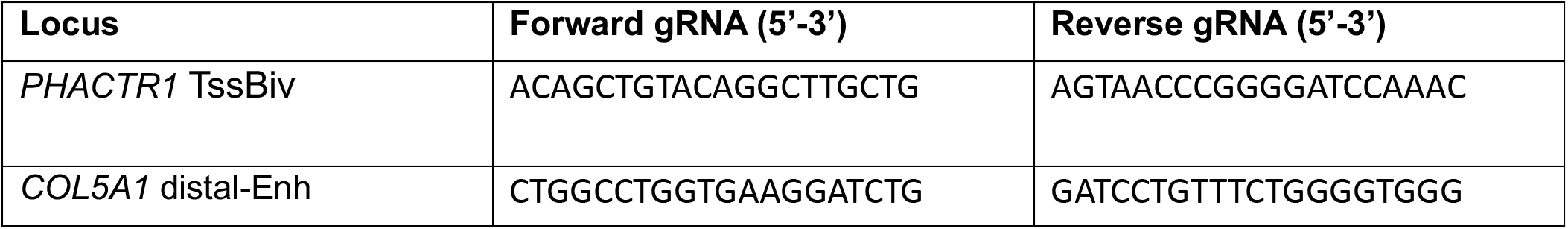
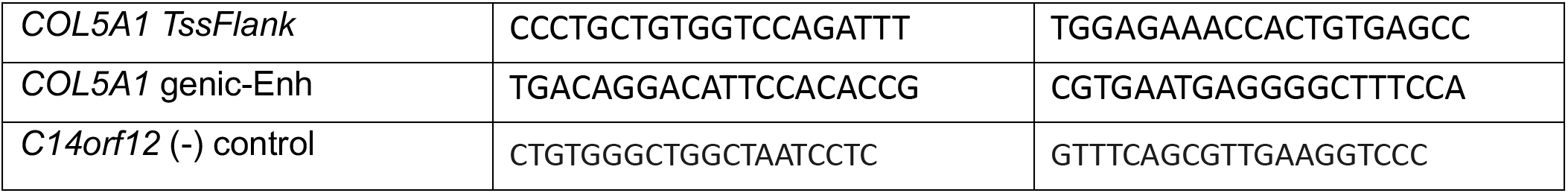

### siQ-ChIP library preparation and sequencing

Immunoprecipitated fragments and saved inputs were quantified with a Qubit dsDNA High Sensitivity Assay kit (Invitrogen, Q32851), and 10 ng of purified DNA for each IP and input sample were used for library preparation with the KAPA Hyper Prep Kit (Kapa Biosystems, KR0961). Library preparation, including fragment end-repair, A-tail extension, and adapter ligation, was conducted per the manufacturer’s instructions (KAPA). Adapter-ligated fragments were amplified with 11 cycles of PCR following the recommended thermocycler program, and DNA was purified with two rounds of purification using KAPA Pure Beads (KK8000). Quality and quantity of prepped libraries were assessed using a combination of Agilent DNA High Sensitivity chip (Agilent Technologies, Inc.), QuantiFluor® dsDNA System (Promega), and Kapa Illumina Library Quantification qPCR assays (Kapa Biosystems). Individually indexed libraries were pooled, and 50 bp paired end sequencing was performed on an Illumina NovaSeq6000 sequencer using an S2 100 bp sequencing kit to a minimum read depth of 50M read pairs per IP library and 100M read pairs per Input library. Base calling was done by Illumina RTA3 and output of NCS was demultiplexed and converted to FastQ format with Illumina Bcl2fastq (v1.9.0).

### siQ-ChIP-seq processing and analysis

siQ-ChIP sequencing reads were 3’ trimmed and filtered for quality and adapter content using TrimGalore (v0.5.0) and quality was assessed by FastQC (v0.11.8). Reads were aligned to human assembly hg38 with bowtie2 (v2.3.5, RRID:SCR_016368) and were deduplicated using removeDups from samblaster (v.0.1.24) (60). Aligned BAM files were used for quality control analysis with “deeptools” (v3.5.4), ‘plotFingerprint,’ and ‘plotPCA’ functions. Aligned SAM files were then processed for pair-end reads with high mapping quality (MAQ ≥ 20), correct pair orientation (Sam Flags = 99, 163), and fragment length as described for siQ-ChIP (https://github.com/BradleyDickson/siQ-ChIP). Param.in files were prepared for each sample with all required parameters and measurements required for siQ-ChIP normalization. IP tracks (with siQ-ChIP efficiency values) and comparative responses between drug treatments (relative to VEH) were generated with execution of getsiq.sh (version: September 2024) with the EXPlayout file (NOTE: params.in files are provided with the GEO accession). Each individual inhibitor-treated biological replicate was compared to each individual vehicle-treated biological replicate. To determine changes in H3K27ac distributions between inhibitor-treated samples and vehicle-treated samples, each inhibitor-treated biological replicate (e.g., DAC1) was individually compared to VEH-treated biological replicates (e.g., DAC1 vs. Veh1, DAC1 vs. Veh2, etc.), and the average log_2_ fold-change in response (area of peak in inhibitor treatment/area of peak in vehicle treatment) was calculated. Peaks were considered conserved among biological replicates using the GenomicRanges (v1.56.2) ‘findOverlaps’ function. Finally, average log_2_ fold-changes in responses were calculated for peaks conserved between the two inhibitor-treated biological replicates. For H3K27ac, peaks were considered significant if the log_2_ fold-change in response was ≥ 0.585 (increase) or ≤ -1.0 (decrease). For H3K27me1, the variance (in general high among the 3 biological replicates) (≤ 1) was considered for significance peak calling along with log_2_ fold-change ≤ -0.5 (decrease) or ≥ 0.5 (increase).

### ChromHMM (v1.23)

Enrichment overlap analysis with siQ-ChIP-quantified peaks were conducted with the ‘OverlapEnrichment’ function using the genomic coordinates for each dataset and the publicly available 18-state chromHMM annotation for parental RKO cells (ENCODE ENCSR974TXE) (61). Enrichment bias for chromHMM states across individual sample analyses were calculated as the percent enrichment for a particular state divided by the total enrichment across all chromHMM states. 18-state chromHMM annotations were condensed into general genomic annotation categories as follows: Promoter (TssA, TssFlnk, TssFlnkD, TssFlnkU, TssBiv), Enhancer (EnhBiv, EnhA1, EnhA2, EnhWk, EnhG1, EnhG2), Genic (TxWk, Tx), and Repressive (ReprPCWk, ReprPC, Het, Quies, ZNF/Rpts).

### Enzymatic-Methyl-seq (EM-seq) library preparation and sequencing

Libraries were prepared by the Van Andel Institute Genomics Core (RRID:SCR_022913) from an input of 41 to 51 ng of ChIP DNA (taken directly from DNA immunoprecipitated for siQ-ChIP-seq) using the NEBNext EM-seq Kit (New England Biolabs, E7120L). The denaturation method used was 0.1 N of sodium hydroxide, according to the protocol, and 8 cycles of PCR amplification were performed. Quality and quantity of the finished libraries were assessed using a combination of the Agilent High Sensitivity DNA chip (Agilent Technologies Inc., 5067-4626) and QuantiFluor dsDNA System (Promega, E2670). Paired-end sequencing (150 bp) was performed on an Illumina NovaSeq6000 sequencer using an S4, 300-bp sequencing kit (Illumina Inc.), with 10% PhiX to a minimum read depth of 100 M read pairs per library. Base calling was done by Illumina RTA3, and output of NCS was demultiplexed and converted to FastQ format with Illumina Bcl2fastq (v1.9.0).

### EM-seq processing and analysis

EM-seq reads were aligned to human genome build hg38, duplicate-marked (samblaster, RRID:SCR_000468), and sorted (samtools, RRID:SCR_002105) using the “biscuitBlaster” pipeline from BISCUIT (v1.7.1) (https://huishenlab.github.io/biscuit/). Cytosine retention and callable single-nucleotide polymorphism mutations were computed with “biscuit pileup” and output into a VCF file. Cytosine 5mC-values and coverage were extracted with “biscuit vcf2bed” and CpG methylation status was merged with “biscuit mergecg.” CpH dinucleotides (control for EM conversion) showed less than 0.5% cytosine methylation indicating that we achieved >99% conversion efficiency on all EM-seq libraries.

CpGs covered by at least 3 reads were retained in downstream analysis for each individual acEM-seq and me1EM-seq sample library. 5mC-values for individual CpGs were converted to bigwig format using UCSC tools (v2.1) (62). Integrated siQ-ChIP-seq and EM-seq analysis was conducted with deeptools (v3.5.4) (63) by constructing matrices with ‘computeMatrix’ across queried genomic coordinates with the respective bigwig data and visualizing the summarized integration with ‘plotProfile’ and ‘plotHeatmap’.

### Genomic DNA isolation

Genomic DNA was extracted using the DNeasy Blood & Tissue Kit (Qiagen, 69504) following the standard protocol. Samples were then treated with 1 mg/ml RNAse A at 37°C for 30 minutes. DNA was re-precipitated with 1/10 volume 3 M sodium acetate pH 4.8 and 2.5 volumes 100% ethanol and stored overnight at -20°C. Precipitated DNA was pelleted by centrifugation at 17,090 g for 30 minutes at 4°C. The pelleted DNA was washed twice with 70% ethanol, allowed to dry for 15 minutes, and resuspended in nuclease-free water.

### Infinium MethylationEPIC BeadChip (EPIC array)

Genomic DNA was quantified with the Qubit dsDNA High Sensitivity Assay kit (Invitrogen, Q32851), and 1.5 µg of genomic DNA was submitted to the VAI Genomics Core (RRID:SCR_022913) for quality control analysis, bisulfite conversion, and DNA methylation quantification using the Infinium MethylationEPIC BeadChIP (Illumina) processed on an Illumina iScan system following the manufacturer’s standard protocol (64,65).

### EPIC array data processing and analysis

All analyses were conducted in the R statistical software (v4.4.1) (R Core Team). Raw IDAT files for each sample were processed using the Bioconductor (RRID:SCR_006442) package “SeSAMe” (version 1.22.2) for extraction of probe signal intensity values, normalization of probe signal intensity values, and calculation of β-values from the normalized probe signal intensity values (66). The β-value is the measure of DNA methylation for each individual CpG probe, where a minimum value of 0 indicates a fully unmethylated CpG and a maximum value of 1 indicates a fully methylated CpG in the population. CpG probes with a detection p-value > 0.05 in any one sample were excluded from the analysis. Differentially methylated regions (DMRs) were called using the Bioconductor package “DMRcate” (v3.0.10), and regions were considered differentially methylated if at least five contiguous CpGs demonstrated a mean difference of 0.20 methylation change in drug-treated cells compared to vehicle (VEH)-treated cells.

### Construction and sequencing of directional total RNA-seq libraries

Libraries were prepared by the Van Andel Institute Genomics Core (RRID:SCR_022913) from 500 ng of total RNA using the KAPA RNA HyperPrep Kit (Kapa Biosystems). Ribosomal RNA material was reduced using the QIAseq FastSelect –rRNA HMR Kit (Qiagen). RNA was sheared to 300-400 bp. Prior to PCR amplification, cDNA fragments were ligated to IDT for Illumina TruSeq UD Indexed adapters (Illumina Inc). Quality and quantity of the finished libraries were assessed using a combination of Agilent DNA High Sensitivity chip (Agilent Technologies, Inc.) and QuantiFluor® dsDNA System (Promega). Individually indexed libraries were pooled, and 50 bp paired end sequencing was performed on an Illumina NovaSeq6000 sequencer to an average depth of 50M raw paired-reads per transcriptome. Base calling was done by Illumina RTA3 and output of NCS was demultiplexed and converted to FastQ format with Illumina Bcl2fastq (v1.9.0).

### RNA-seq processing and analysis

Raw 50 bp paired-end reads were trimmed with TrimGalore! (http://www.bioinformatics.babraham.ac.uk/projects/trim_galore/) (RRID:SCR_011847) followed by quality control analysis with FastQC. Trimmed reads were aligned to GRCh38.p12 and indexed to GENCODE v29 via STAR (v2.5.3a) aligner with flags ‘-twopassMode Basic\ -quantMode GeneCounts’ for feature counting.

ReadsPerGene output count files were constructed into a raw read count matrix in R. Low count genes were filtered (1 count in at least one sample) prior to edgeR (v4.2.2) count normalization and differential expression analysis with voomWithQualityWeights and quasi-likelihood fit set to robust. Principal component analysis was calculated using ‘prcomp’ in the R stats package on the normalized expression matrix. Differential expression analysis was conducted with voomWithQualityWeights and a quasi-likelihood fit set to robust. Each dataset was treated separately for differential expression analysis, and all inhibitor treatments were compared to their respective vehicle samples. Genes were considered differentially expressed if |log_2_FC| ≥ 1 and FDR ≤ 0.01.

### Gene Set Enrichment Analysis (GSEA)

GSEA (v4.3.3) (67) was conducted across the HALLMARK and c2.cp.kegg curated gene set databases from Molecular Signature Database (MSigDB). Phenotype comparisons were set to the sample of interest (e.g., VD) VS REST (monotherapy treatments; e.g., VEH/DAC/VAL) for all analyses with weighted enrichment statistic and Signal2Noise settings for ranking genes. Maximum and minimum size of a gene set was defined as 1500 and 15 genes, respectively. Genes marked as a “core enrichment” gene were used for epigenetic profiling and heatmap row z-score analysis of normalized cpm values.

### Assay for Transposase-Accessible Chromatin (ATAC) sequencing

ATAC sequencing was performed using an ATAC-Seq Kit (ActiveMotif, 53156) according to the manufacturer’s instructions with specific modifications. Briefly, RKO cells from each treatment group were trypsinized, harvested, and processed immediately. For each treatment, 50,000 cells were incubated with Tn5 transposase for 45 min to optimize nucleosomal laddering. Following PCR amplification of the tagmented DNA, a right-sided Solid Phase Reversible Immobilization (SPRI) bead cleanup was performed to remove high-molecular-weight DNA fragments. Briefly, 0.5× volume of SPRI beads (provided with the kit) was added to each sample, vortexed briefly, and incubated for 5 min at room temperature. Samples were then placed on a magnetic stand, and the supernatant containing the desired smaller fragments was transferred to a fresh tube. This supernatant was subsequently processed through the standard SPRI bead purification protocol as recommended by the manufacturer. The final purified libraries were submitted for high-throughput sequencing.

Libraries were sequenced by the Van Andel Genomics Core. Quality and quantity of the libraries were assessed using a combination of Agilent TapeStation D5000 ScreenTape analysis (Agilent Technologies, Inc.), and QuantiFluor® dsDNA System (Promega Corp., Madison, WI, USA). Individually indexed libraries were pooled, and 75 bp paired-end sequencing was performed on an Element AVITI sequencer using a high-output, 75 bp sequencing kit (Element Biosciences, San Diego, CA, USA) to an average depth of 50M reads. Base calling was performed on instrument and AVITI OS (v2.6.2) output was demultiplexed and converted to FastQ format with Element Biosciences Bases2fastq (v1.7.0).

### ATAC-seq pre-processing and analysis

Paired-end ATAC-seq reads were adapter- and quality-trimmed with Trim Galore (v0.6.10) using Cutadapt (v2.6) (‘--quality 25 –paired’), with FastQC performed on the trimmed reads. Trimmed reads were aligned to the hg38 genome with Bowtie2 using the ‘--very-sensitive’, ‘-I 0’, and ‘-X 1000’ settings. Duplicate fragments were removed with SAMBLASTER (‘—removeDups’). Properly paired, primary alignments with mapping quality at least 30 were retained using SAMtools v1.21 (‘samtools view -b -q 30 -f 2 -F 2828’), and alignments to chrM were removed.

Peaks were called separately for each sample with MACS3 (v3.0.3) in paired-end mode (‘callpeak --format BAMPE --gsize hs --qvalue 0.01 --keep-dup all --bdg –SPMR’). The ‘--call-summits’ option was not used. Peak intervals overlapping the hg38 blacklist or non-canonical chromosomes were removed. Within each treatment, genomic segments supported by peaks in at least two of the three biological replicates were retained and adjacent supported segments were merged. Fragments overlapping consensus intervals were quantified with featureCounts (v2.1.1) in paired-end, fragment-counting mode using a SAF annotation (‘-F SAF -p --countReadPairs -B -C --primary -Q 30 -s 0’).

The consensus count matrix contained 18 RKO samples representing VEH, TAZ, VAL, DAC, TD, and VD (n=3 per condition). Consensus peaks were retained when at least 10 fragments were detected in at least three samples. Counts were analyzed with DESeq2 (v3.23) using the design ‘∼ block + condition’, with VEH as the reference condition. Variance-stabilized counts were used for principal-component and sample-correlation analyses.

Differential accessibility was calculated for TAZ, VAL, DAC, TD, and VD relative to VEH, as well as for the combination-versus-single-agent contrasts. Peaks were classified as significantly increased at a Benjamini-Hochberg-adjusted P-value <= 0.05 and log_2_FC >= 1, and significantly decreased at an adjusted P-value <= 0.05 and log_2_ FC <= -1.

Significantly increased TD and VD peaks were compared using their identifiers in the common consensus peak universe. Peaks significant only under VD were designated VD-only. Peaks significant under both TD and VD were divided using the prespecified magnitude rule: VD log_2_FC >= 1.20 x TD log_2_FC. Shared peaks satisfying this rule were classified as at least 20% stronger under VD; the remaining shared peaks were classified as “Comparable” TD/VD responses. For downstream analyses, VD-only peaks and shared peaks at least 20% stronger under VD were combined into the VD-favored class. This produced 10,020 VD-favored increased-accessibility peaks and 5,654 comparable TD/VD peaks. Classification was performed before incorporating RNA-seq data.

Edge-to-edge distances between accessibility peaks and differential H3K27me1 intervals were calculated with ‘distanceToNearest’ from GenomicRanges while ignoring strand. Peaks within 10 kb of TAZ-associated H3K27me1 gain or TAZ-maintained/VAL-only H3K27me1 loss were retained for the integrated analysis. To create mutually exclusive assignments, H3K27me1 gain was prioritized over maintenance when a peak was within 10 kb of both classes. Peaks without an H3K27me1 interval within 10 kb were not included in the chromatin-barrier analysis. The combinations were interpreted as follows with regard to assigned chromatin barrier classification: VD-favored accessibility near TAZ-induced H3K27me1 gain, adaptive H3K27me1 barrier; Comparable accessibility near H3K27me1 gain or VD-favored accessibility near TAZ-maintained H3K27me1, H3K27me1 plus induced bivalency; and Comparable accessibility near TAZ-maintained H3K27me1, induced bivalency.

Accessibility peaks were associated with genes using rGREAT and the hg38 GREAT transcription-start-site (TSS) annotation. The basal-plus-extension rule was used with basal domains extending 5 kb upstream and 1 kb downstream of each TSS and extension up to 1 Mb or the basal domain of the neighboring gene. The default whole-genome background excluding assembly gaps was used. BED zero-based starts were converted to one-based GRanges coordinates before analysis. Peak-gene links within the -5-kb to +1-kb GREAT basal domain were classified as TSS; extended-domain links were classified as candidate cis-regulatory elements (CREs).

GREAT-associated genes were joined to the annotated RNA-seq differential-expression table by gene symbol. Symbols represented by more than one RNA-seq record were considered ambiguous and excluded. To prevent RNA results from influencing peak selection, one representative peak was selected per gene using the following expression-blind hierarchy: TSS over CRE; H3K27me1 gain over maintenance and loss; VD-favored over Comparable accessibility; shortest absolute GREAT distance to the TSS; and peak identifier as the final deterministic tie-breaker.

### Integrative genomic analysis

As described in the respective sections above, bigwig files were generated for each sample for genome-wide H3K27ac and H3K27me1 efficiency (siQ-ChIP-seq), DNA methylation β-values (EPIC array), and publicly available datasets for RKO parental Whole Genome Bisulfite Sequencing (GSE262054, WGBS) (68) and our previously published H3K27me3 siQ-ChIP data (GSE256135) for parental- and DNMT1i- (GSK3484862, 1 µM) treated RKO cells (12). Bed files with genomic coordinates for differential H3K27 peak analysis and ChromHMM chromatin state annotations were generated as described above. Integrated siQ-ChIP-seq, WGBS, and EPIC array analysis was conducted with deeptools (v3.5.4) (63) by constructing matrices with ‘computeMatrix’ across queried genomic coordinates with the respective bigwig data and visualizing the summarized integration with ‘plotProfile’ and ‘plotHeatmap’. Motif enrichment analysis was conducted using ‘findMotifsGenome.pl -len 8,10,12’ from HOMER (v4.11.1) (69).

