## Supplemental Figures for "Combined EZH1/2 and DNMT inhibition overcomes transcriptional barriers imposed by drug-induced epigenetic plasticity in colorectal cancer cells"

Fig S1

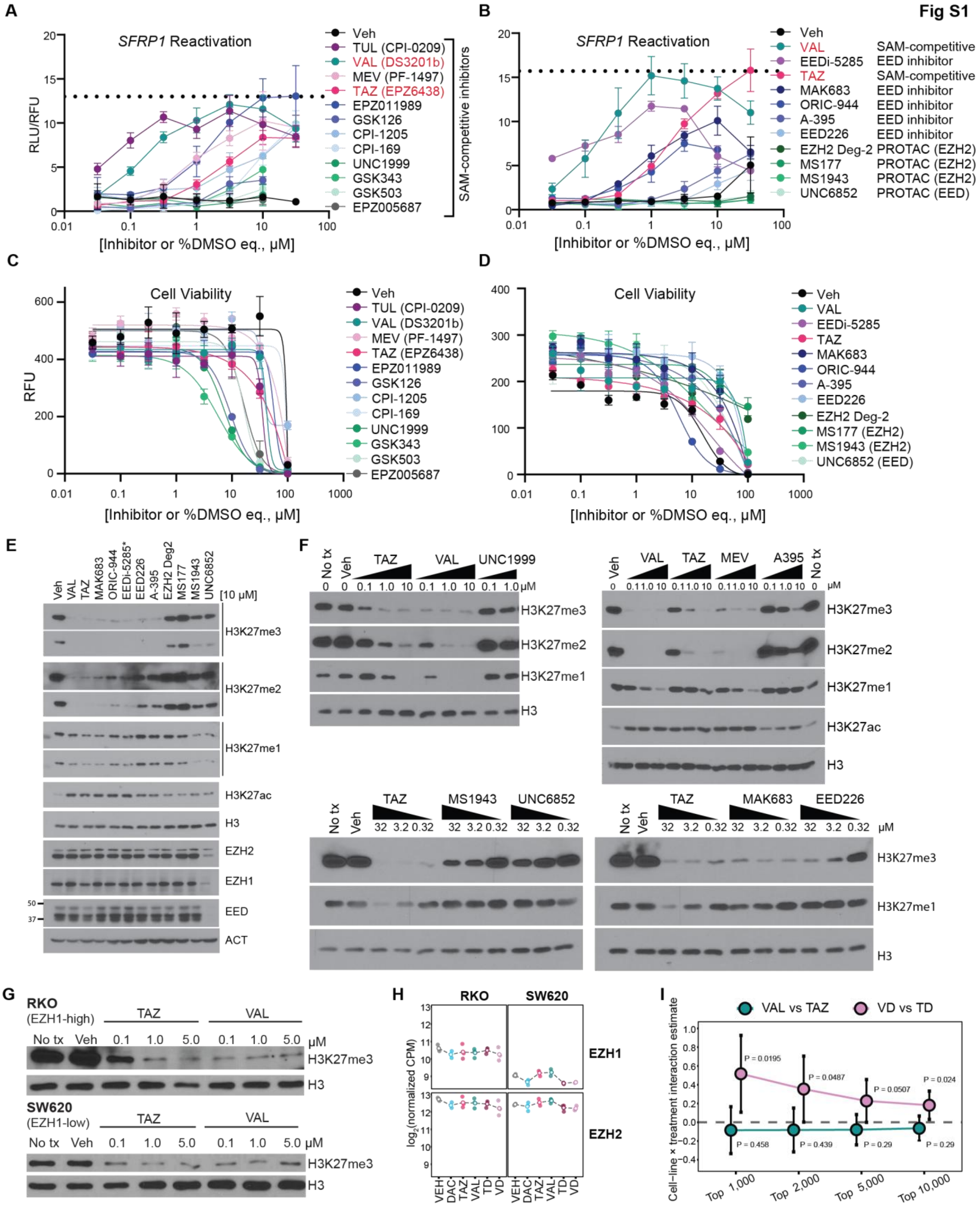

A
SFRP1 Reactivation
RLU/RFU
[Inhibitor or %DMSO eq., μM]
Veh
TUL (CPI-0209)
VAL (DS3201b)
MEV (PF-1497)
TAZ (EPZ6438)
EPZ011989
GSK126
CPI-1205
CPI-169
UNC1999
GSK343
GSK503
EPZ005687
SAM-competitive inhibitors
B
SFRP1 Reactivation
Veh
VAL SAM-competitive
EEDi-5285 EED inhibitor
TAZ SAM-competitive
MAK683 EED inhibitor
ORIC-944 EED inhibitor
A-395 EED inhibitor
EED226 EED inhibitor
EZH2 Deg-2 PROTAC (EZH2)
MS177 PROTAC (EZH2)
MS1943 PROTAC (EZH2)
UNC6852 PROTAC (EED)
[Inhibitor or %DMSO eq., μM]
C
Cell Viability
RFU
[Inhibitor or %DMSO eq., μM]
D
Cell Viability
Veh
VAL
EEDi-5285
TAZ
MAK683
ORIC-944
A-395
EED226
EZH2 Deg-2
MS177 (EZH2)
MS1943 (EZH2)
UNC6852 (EED)
[Inhibitor or %DMSO eq., μM]
E
Veh VAL TAZ MAK683 ORIC-944 EEDi-5285* EED226 A-395 EZH2 Deg2 MS177 MS1943 UNC6852 [10 μM]
H3K27me3
H3K27me2
H3K27me1
H3K27ac
H3
EZH2
EZH1
EED
ACT
F
No tx Veh TAZ VAL UNC1999
0 0 0.1 1.0 10 0.1 1.0 10 0.1 1.0 μM
H3K27me3
H3K27me2
H3K27me1
H3
Veh VAL TAZ MEV A395 No tx
H3K27me3
H3K27me2
H3K27me1
H3K27ac
H3
No tx Veh TAZ MS1943 UNC6852
32 3.2 0.32 32 3.2 0.32 32 3.2 0.32
No tx Veh TAZ MAK683 EED226
32 3.2 0.32 32 3.2 0.32 32 3.2 0.32 μM
H3K27me3
H3K27me1
H3
G
RKO
(EZH1-high)
No tx Veh TAZ VAL
0.1 1.0 5.0 0.1 1.0 5.0 μM
H3K27me3
H3
SW620
(EZH1-low)
No tx Veh TAZ VAL
0.1 1.0 5.0 0.1 1.0 5.0 μM
H3K27me3
H3
H
RKO SW620
log2(normalized CPM)
EZH1
EZH2
VEH DAC TAZ VAL TD VD
I
VAL vs TAZ
VD vs TD
Cell-line × treatment interaction estimate
P = 0.0195
P = 0.0487
P = 0.0507
P = 0.024
P = 0.458
P = 0.439
P = 0.29
P = 0.29
Top 1,000
Top 2,000
Top 5,000
Top 10,000

**Figure S1. Dual EZH1/2 inhibitors block all three H3K27 methylation states**

**A-B**) NLuc reporter activity measurements following 72-hour treatment with the indicated PRC2 antagonists. Red text indicates SAM-competitive inhibitors shared between experiments. Relative luminescence units (RLU) are normalized to relative fluorescence units (RFU) from multiplexed CellTiter-Fluor viability assays. Data are mean ± SD of technical triplicates and are representative of n=3 biological replicates. Dotted black lines denote the maximal signal measured without the addition of DAC. TUL, tulmimetostat; VAL, valemetostat; MEV, mevrometostat; and TAZ, tazemetostat.

**C-D**) CellTiter-Fluor dose-response viability curves for **A-B**.

**E**) Western blot analysis of the indicated proteins and PTMs from wild-type HCT116 cells following 72-hour treatment with vehicle (DMSO, % equivalent) or the indicated PRC2 inhibitors (10 µM) ordered approximately by ability to induce SFRP1 expression as reported in **Figure 1B**.

**F**) Western blot analysis of all H3K27 methylation states from wild-type HCT116 cells following 72-hour treatment with vehicle (DMSO, % equivalent) or titrations of the indicated PRC2 inhibitors.

**G)** Western blot analysis of H3K27me3 states from wild-type RKO cells following 72-hour treatment with vehicle (DMSO, % equivalent) or titrations of the indicated PRC2 inhibitors.

**H)** EZH1 and EZH2 transcript abundance in RKO and SW620 cells following treatment with vehicle (VEH), decitabine (DAC), tazemetostat (TAZ), valemetostat (VAL), DAC plus TAZ (TD), or DAC plus VAL (VD). Points represent biological replicates (n=3); open circles indicate treatment means, and lines connect means.

**I)** Cell-line x treatment interaction estimate differences in RNA expression between VD vs TD or VAL vs TAZ treatment in RKO versus SW620 cells. P-values indicate significance between cell line responses for the indicated drug treatments.

**Related to Figure 1.**

**Fig S2**

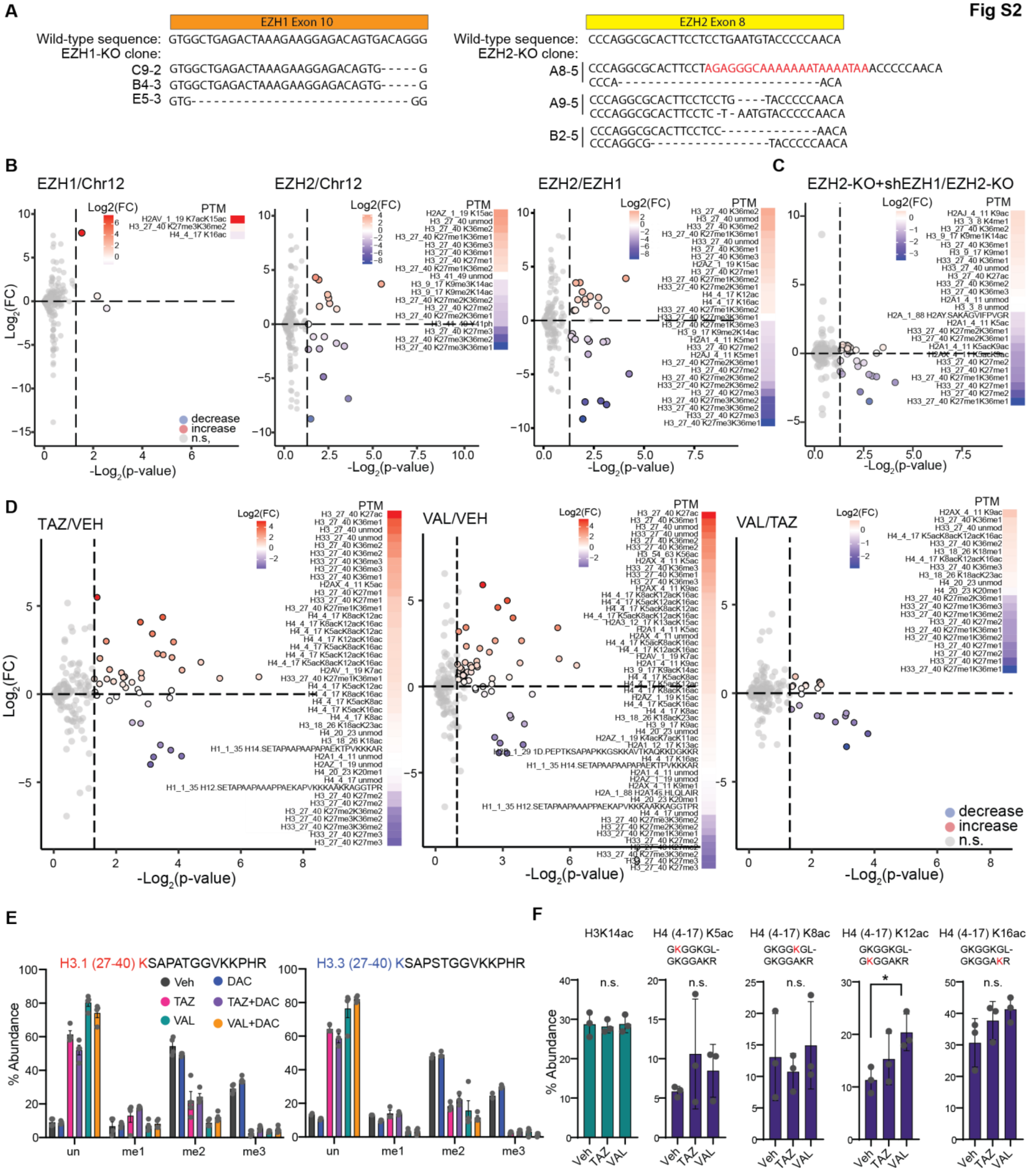
A
EZH1 Exon 10
Wild-type sequence: GTGGCTGAGACTAAAGAAGGAGACAGTGACAGGG
EZH1-KO clone:
C9-2 GTGGCTGAGACTAAAGAAGGAGACAGTG-----G
B4-3 GTGGCTGAGACTAAAGAAGGAGACAGTG-----G
E5-3 GTG------------------------------GG
EZH2 Exon 8
Wild-type sequence: CCCAGGCGCACTTCCTCCTGAATGTACCCCCAACA
EZH2-KO clone:
A8-5 CCCAGGCGCACTTCCTAGAGGGCAAAAAAATAAAATAAACCCCCAACA
CCCA--------------------------ACA
A9-5 CCCAGGCGCACTTCCTCCTG----TACCCCCAACA
CCCAGGCGCACTTCCTC-T-AATGTACCCCCAACA
B2-5 CCCAGGCGCACTTCCTCC-------------AACA
CCCAGGCG----------------TACCCCCAACA
B
EZH1/Chr12
EZH2/Chr12
EZH2/EZH1
C
EZH2-KO+shEZH1/EZH2-KO
Log2(FC)
PTM
Log$_2$(FC)
-Log$_2$(p-value)
decrease
increase
n.s.
D
TAZ/VEH
VAL/VEH
VAL/TAZ
E
H3.1 (27-40) KSAPATGGVKKPHR
H3.3 (27-40) KSAPSTGGVKKPHR
Veh
DAC
TAZ
TAZ+DAC
VAL
VAL+DAC
% Abundance
un
me1
me2
me3
F
H3K14ac
H4 (4-17) K5ac
GKGGKGL-GKGGAKR
H4 (4-17) K8ac
H4 (4-17) K12ac
H4 (4-17) K16ac
n.s.
*
Veh
TAZ
VAL

**Figure S2. Proteomics analysis of histone methylation states following EZH1/2 loss or inhibition.**
**A)** CRISPR/Cas9 indel sequencing of EZH1- and EZH2-targeted RKO clones shown in Figure 3A. Red indicates insertion; dashes indicate deletions. Note: EZH1 clones showed homozygous indels for both alleles.
**B)** Volcano plots of differential histone PTMs in comparisons among EZH1-KO, EZH2-KO, and Chr12 control clones. Red indicates significantly increased PTMs; blue indicates significant decreased PTMs; gray indicates no significant change. Fold changes (FC) were calculated from average abundances of three biological replicate clones per genotype.
**C)** Volcano plots of differential histone PTMs in EZH2-KO cells following shEZH1 induction relative to EZH2-KO alone. FC was calculated from average abundances of n=3 biological replicates.
**D)** Volcano plots of differential histone PTMs in comparisons among RKO cells treated for 72 hours with TAZ (1 µM), VAL (1 µM), or vehicle (DMSO, % equivalent). FC was calculated from average abundances of n=3 biological replicates.
**E-F)** Histone PTM mass spectrometry analysis of relative abundances of H3K27 methylation states on H3.1 and H3.3 (**E**) or the indicated H3 acetylations (**F**) in RKO cells treated for 72 hours with TAZ (1 µM), VAL (1 µM), DAC (300 nM), or the indicated combinations. Data are mean ± SD of biological triplicates. Red or blue "K" denotes modified lysine residues. Statistical significance was calculated using multiple unpaired t-tests. ns, not significant. *$p<0.05$, **$p<0.01$, ***$p<0.001$.
**Related to Figure 2.**

Fig S3

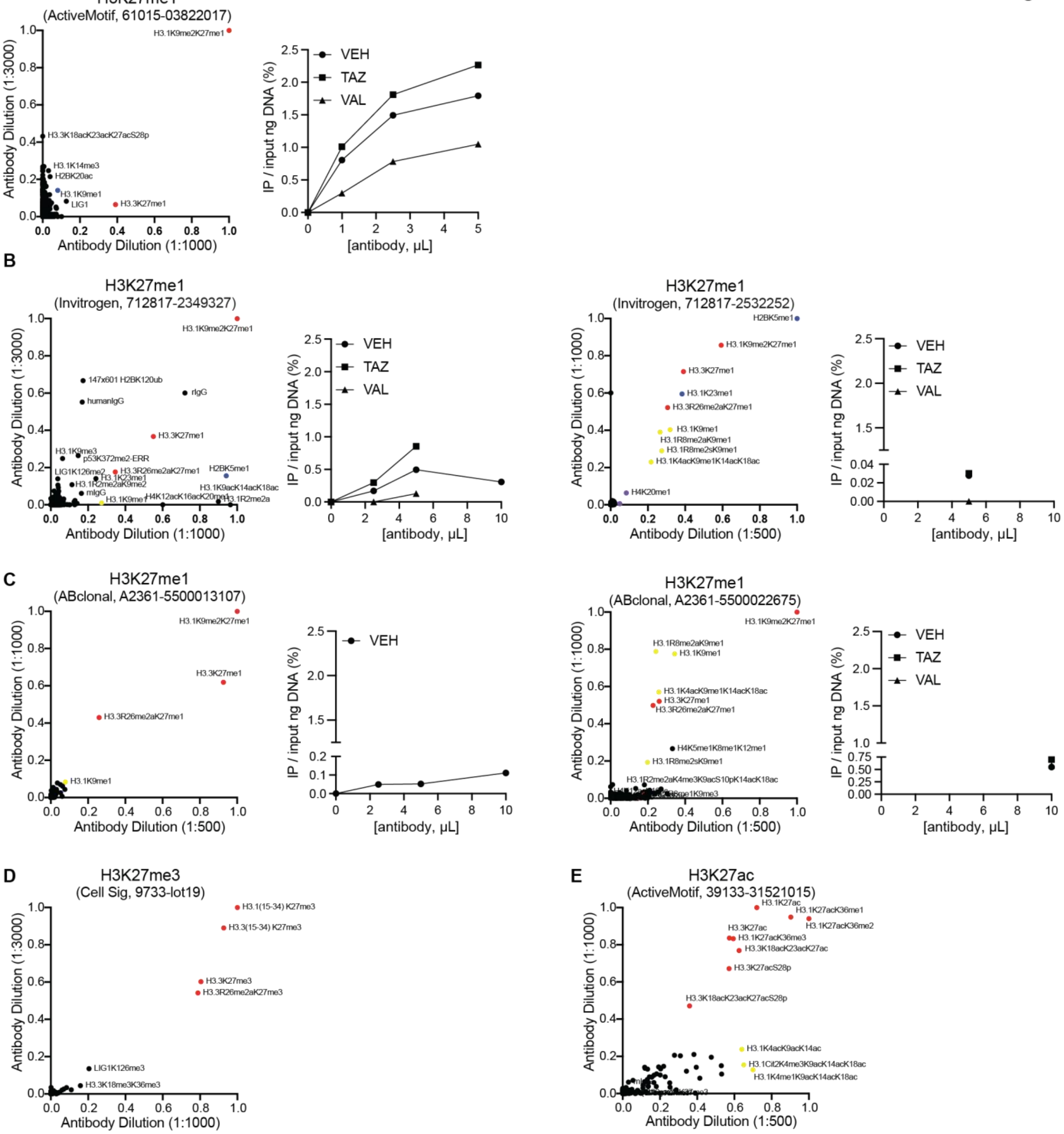
A
H3K27me1
(ActiveMotif, 61015-03822017)
Antibody Dilution (1:3000)
Antibody Dilution (1:1000)
H3.1K9me2K27me1
H3.3K18acK23acK27acS28p
H3.1K14me3
H2BK20ac
H3.1K9me1
LIG1
H3.3K27me1
IP / input ng DNA (%)
[antibody, μL]
VEH
TAZ
VAL
B
H3K27me1
(Invitrogen, 712817-2349327)
Antibody Dilution (1:3000)
Antibody Dilution (1:1000)
H3.1K9me2K27me1
147x601 H2BK120ub
rIgG
humanIgG
H3.3K27me1
H3.1K9me3
p53K372me2-ERR
H2BK5me1
LIG1K126me2
H3.3R26me2aK27me1
H3.1K23me1
H3.1R2me2aK9me2
H3.1K9acK14acK18ac
mIgG
H3.1K9me1
H4K12acK16acK20me1
H3.1R2me2a
IP / input ng DNA (%)
[antibody, μL]
VEH
TAZ
VAL
H3K27me1
(Invitrogen, 712817-2532252)
Antibody Dilution (1:1000)
Antibody Dilution (1:500)
H2BK5me1
H3.1K9me2K27me1
H3.3K27me1
H3.1K23me1
H3.3R26me2aK27me1
H3.1K9me1
H3.1R8me2aK9me1
H3.1R8me2sK9me1
H3.1K4acK9me1K14acK18ac
H4K20me1
IP / input ng DNA (%)
[antibody, μL]
VEH
TAZ
VAL
C
H3K27me1
(ABclonal, A2361-5500013107)
Antibody Dilution (1:1000)
Antibody Dilution (1:500)
H3.1K9me2K27me1
H3.3K27me1
H3.3R26me2aK27me1
H3.1K9me1
IP / input ng DNA (%)
[antibody, μL]
VEH
H3K27me1
(ABclonal, A2361-5500022675)
Antibody Dilution (1:1000)
Antibody Dilution (1:500)
H3.1K9me2K27me1
H3.1R8me2aK9me1
H3.1K9me1
H3.1K4acK9me1K14acK18ac
H3.3K27me1
H3.3R26me2aK27me1
H4K5me1K8me1K12me1
H3.1R8me2sK9me1
H3.1R2me2aK4me3K9acS10pK14acK18ac
IP / input ng DNA (%)
[antibody, μL]
VEH
TAZ
VAL
D
H3K27me3
(Cell Sig, 9733-lot19)
Antibody Dilution (1:3000)
Antibody Dilution (1:1000)
H3.1(15-34) K27me3
H3.3(15-34) K27me3
H3.3K27me3
H3.3R26me2aK27me3
LIG1K126me3
H3.3K18me3K36me3
E
H3K27ac
(ActiveMotif, 39133-31521015)
Antibody Dilution (1:1000)
Antibody Dilution (1:500)
H3.1K27ac
H3.1K27acK36me1
H3.1K27acK36me2
H3.3K27ac
H3.1K27acK36me3
H3.3K18acK23acK27ac
H3.3K27acS28p
H3.3K18acK23acK27acS28p
H3.1K4acK9acK14ac
H3.1Cit2K4me3K9acK14acK18ac
H3.1K4me1K9acK14acK18ac

**Figure S3. Validation of ChIP antibody specificity and sensitivity.**

**A-E)** Antibody validation using histone peptide microarrays (left panels) and ChIP titrations of chromatin isolated from RKO cells (right panels). DNA recovery is expressed as % input across antibody concentrations. Antibodies evaluated: H3K27me1 (Active Motif, **A**; Invitrogen, **B**; ABClonal, **C**), H3K27me3 (Cell Signaling, **D**), and H3K27ac (ActiveMotif, **E**).

**F)** Summary of antibodies and array specificity.

**Related to Figure 3.**

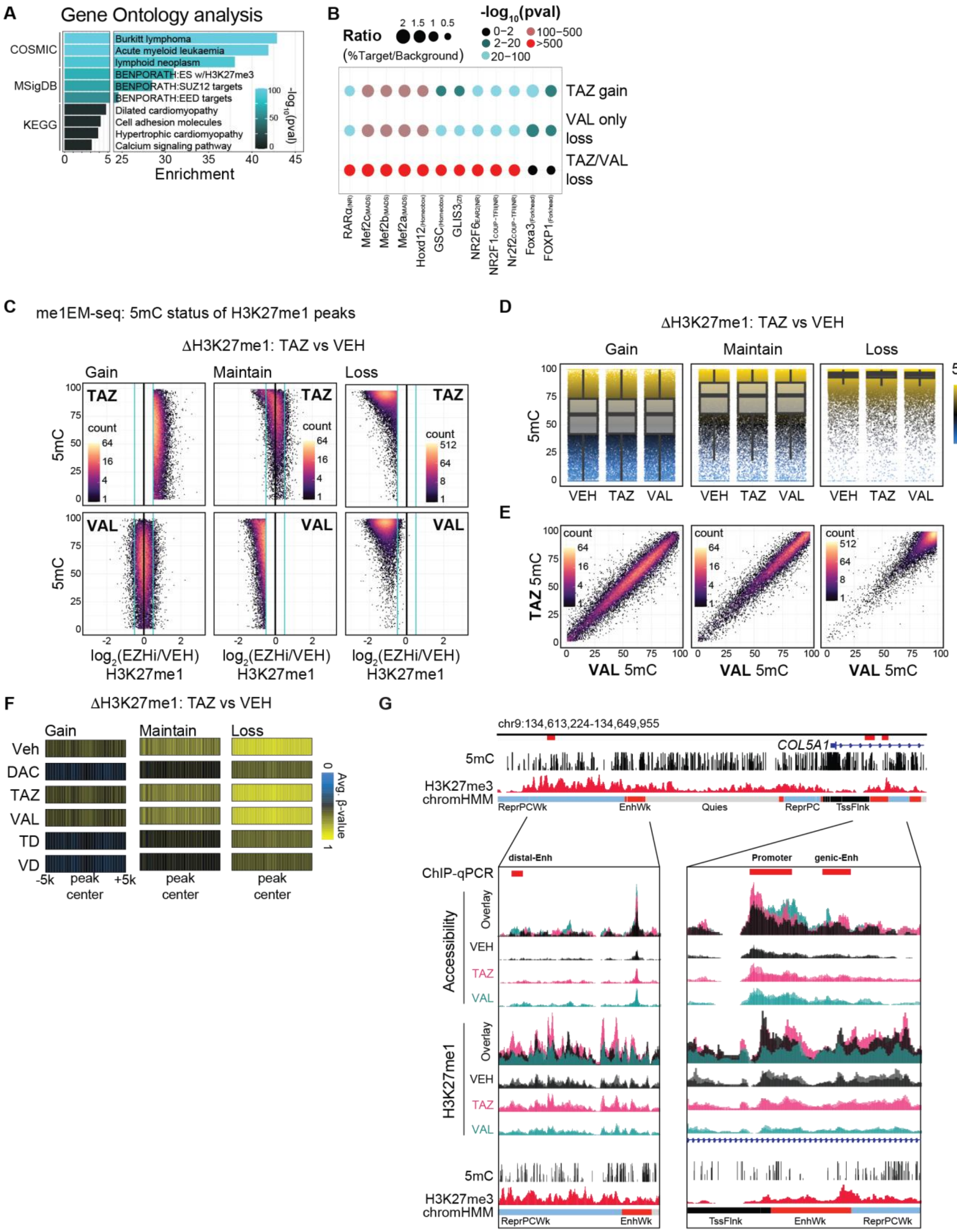
A
Gene Ontology analysis
COSMIC
MSigDB
KEGG
Burkitt lymphoma
Acute myeloid leukaemia
lymphoid neoplasm
BENPORATH:ES w/H3K27me3
BENPORATH:SUZ12 targets
BENPORATH:EED targets
Dilated cardiomyopathy
Cell adhesion molecules
Hypertrophic cardiomyopathy
Calcium signaling pathway
Enrichment
-log10(pval)
B
Ratio (%Target/Background)
-log10(pval)
0-2
2-20
20-100
100-500
>500
TAZ gain
VAL only loss
TAZ/VAL loss
RARα
Mef2c
Mef2b
Mef2a
Hoxd12
GSC
GLIS3
NR2F6
NR2F1
Nr2f2
Foxa3
FOXP1
C
me1EM-seq: 5mC status of H3K27me1 peaks
ΔH3K27me1: TAZ vs VEH
Gain
Maintain
Loss
TAZ
VAL
count
5mC
log2(EZHi/VEH) H3K27me1
D
ΔH3K27me1: TAZ vs VEH
Gain
Maintain
Loss
5mC
VEH
TAZ
VAL
E
count
TAZ 5mC
VAL 5mC
F
ΔH3K27me1: TAZ vs VEH
Gain
Maintain
Loss
Veh
DAC
TAZ
VAL
TD
VD
-5k
peak center
+5k
Avg. β-value
G
chr9:134,613,224-134,649,955
COL5A1
5mC
H3K27me3
chromHMM
ReprPCWk
EnhWk
Quies
ReprPC
TssFlnk
distal-Enh
Promoter
genic-Enh
ChIP-qPCR
Accessibility
Overlay
VEH
TAZ
VAL
H3K27me1

**Figure S4. Motif enrichment and DNA methylation analyses at regions of H3K27me1 dynamics following EZHi and/or DNMTi**

**A)** Gene ontology analysis for TAZ-specific H3K27me1 gains in Figure 3C-D.
**B)** HOMER motif enrichment analysis for EZHi-specific H3K27me1 gains and losses described in Figure 4C-D. Size of bubble indicates degree of enrichment over background as the ratio of (% H3K27me1 change with motif target) / (% background with motif target); color indicates significance.
**C)** Density scatterplots from H3K27me1 siQ-ChIP coupled to enzymatic methyl sequencing (me1EM-seq) showing 5mC levels within altered H3K27me1 peaks (left: TAZ-specific gains; middle: TAZ maintain; right: TAZ loss).
**D)** Boxplots of me1EM-seq DNA methylation distribution within altered peaks of H3K27me1 across treatment conditions.
**E)** Density scatterplots comparing VAL versus TAZ 5mC levels from me1EM-seq at altered H3K27me1 peaks.
**F)** Average DNA methylation profiles across the indicated drug treatments, centered on altered H3K27me1 peaks (left, TAZ-specific gains; middle, VAL-specific losses; right, shared losses).
**G)** Representative genome browser track for the *COL5A1* locus demonstrating chromatin accessibility and H3K27me1 levels across VEH, TAZ, and VAL treatments. Baseline DNA methylation (5mC) and H3K27me3 are presented with ChromHMM state annotations. Regions tested for H3K27me1 ChIP-qPCR (**Figure 3K**) are indicated.
**Related to Figure 3.**

Fig S5

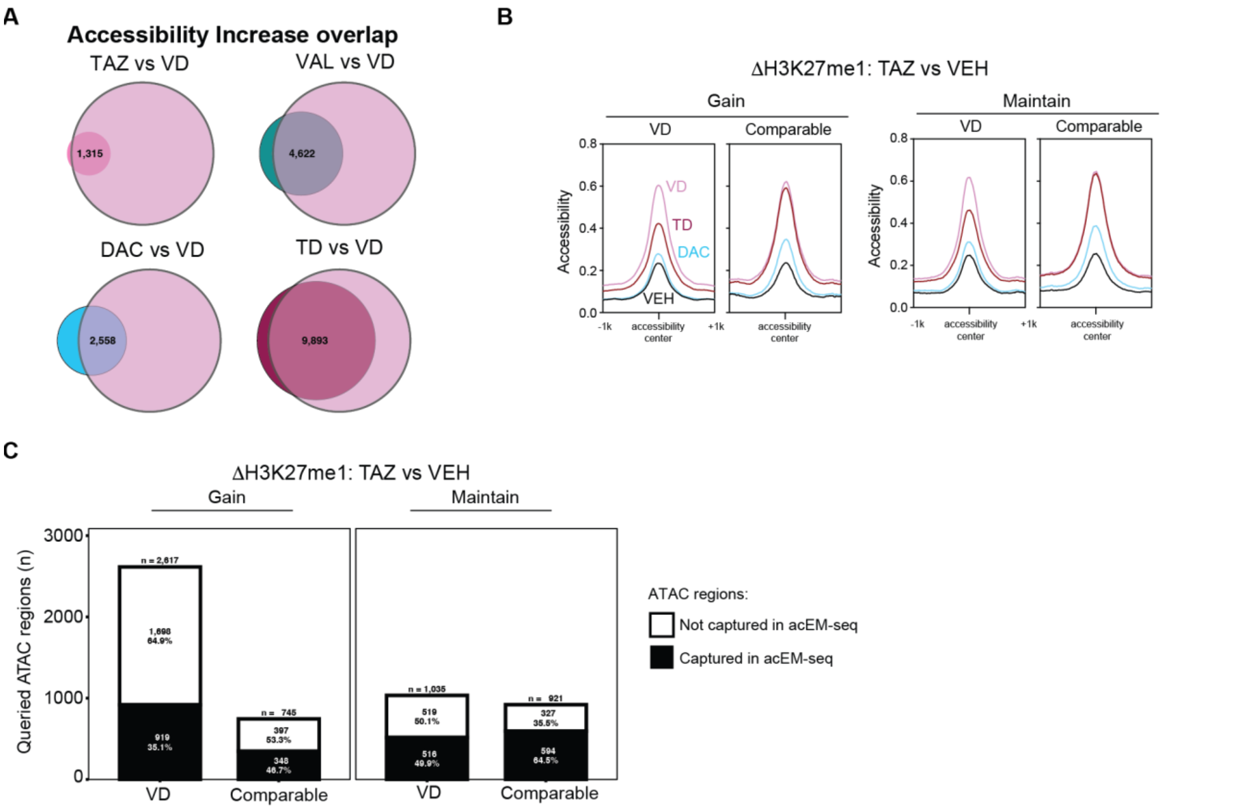


**Figure S5. Chromatin accessibility classification.**
**A)** Venn diagrams of overlapping regions of increased accessibility among the TAZ, VAL, DAC, and TD compared to VD. Number ATAC regions overlapping among treatment paradigms is indicated.
**B)** Average chromatin accessibility across treatments at indicated classified increased accessibility regions (from **Figure 4B**).
**C)** Bar graph of increased accessibility regions (from **Figure 4B**) that were sufficiently captured by acEM-seq for DNA methylation analysis presented in **Figure 4G**.
**Related to Figure 4.**
